# Gene-metabolite networks reveal physiological trade-offs but not transcriptional co-regulation between carotenoid and dry matter accumulation in cassava *(Manihot esculenta)* roots

**DOI:** 10.64898/2026.07.11.737982

**Authors:** Seren S. Villwock, Kiara M. Gómez, Tara Fish, Jiho Lee, Ailene Doherty, Adrian White, Theodore Thannhauser, Michael A. Gore, Jean-Luc Jannink

## Abstract

Provitamin A biofortification of cassava (*Manihot esculenta*) is constrained by a negative genetic correlation between total carotenoid (TC) and dry matter (DM) contents, but its underlying biological mechanism remains unclear. We examined this relationship in 24 African, Latin American, and hybrid cassava genotypes by measuring gene expression and metabolites in inner and outer storage root layers that varied spatially in carotenoid accumulation, enabling comparisons across and within genotypes. TC, and particularly upstream *cis-*carotenes, were negatively associated with DM after accounting for genotype, and this relationship strengthened across developmental timepoints, consistent with a physiological component of the trade-off. Gene-metabolite co-expression networks constructed from genetic, non-genetic, and overall phenotypic trait components showed no significant topological overlap between carotenoid- and starch-related gene sets, indicating that the TC/DM relationship is not driven by transcriptional co-regulation of the main pathways, although individual associations between carotenoids and carbohydrate-related genes suggested possible indirect metabolic links. Instead, TC accumulation was most strongly associated with regulatory, stress-, redox-, and plastid-related genes, enriched for heme-binding and oxidoreductase functions. These findings suggest that the TC/DM trade-off is not a fixed constraint, and identify candidate regulators of carotenoid content for future functional validation to support cassava provitamin A biofortification.

**Highlight:** Genetic, spatial, and developmental variation in cassava storage roots implicate *cis-*carotenes, but not direct transcriptional regulation of the carotenoid and starch pathways, in the trade-off between carotenoids and dry matter.

## Introduction

Micronutrient deficiencies are estimated to affect the health of two billion people globally, particularly in impoverished regions that lack reliable access to dietary diversity (Black et al., 2013; Stevens et al., 2015; Ritchie and Roser, 2017). Nearly half of all young children in sub-Saharan Africa are at risk for vitamin A deficiency, which can cause blindness, immunosuppression, stunting, and mortality (Stevens et al., 2015). Vitamin A deficiency rates are particularly high in communities whose caloric intake relies heavily on cassava (*Manihot esculenta*), a resilient food security crop with starchy storage roots that is a staple food for millions of smallholder farmers in tropical regions (Montagnac et al., 2009; Waisundara, 2018). Though cassava is generally low in micronutrients, some yellow-fleshed cassava varieties naturally accumulate low amounts of provitamin A carotenoids (predominantly the orange-pigmented *β*-carotene), which increases both nutritional value and post-harvest storability (Sánchez et al., 2006; Esuma et al., 2012; Atser, 2014; Beyene et al., 2018; Afolami et al., 2021).

Recent initiatives have focused on the provitamin A biofortification of cassava and other staple crops to help address vitamin A deficiency (Saltzman et al., 2013; Giuliano, 2017; Beyene et al., 2018). However, these efforts have been challenged by a negative genetic correlation between total carotenoid (TC) and dry matter (DM) contents in cassava (reported correlation *r* ranging from -0.59 to -0.3 in African breeding populations), which compromises starch yield, cooking quality, and consumer acceptance (Dixon et al., 2008; Fukuda et al., 2010; Tahirou et al., 2015; Rabbi et al., 2017; Atwijukire et al., 2019). Similar relationships have also been observed in other starchy crops, including sweetpotato, squash, and citrus (Cao et al., 2015; Grisales et al., 2015; Mortimer et al., 2016; Gemenet et al., 2020), reviewed in Villwock et al. (2024).

A major driver of carotenoid accumulation in cassava is a genetic variant in the *phytoene synthase 2 (PSY2)* gene, which encodes the key rate-limiting enzyme in carotenoid biosynthesis. The causal *PSY2-Y-2* allele, a single C_572_A mutation, has been functionally characterized and is known to increase the catalytic efficiency of PSY2 (Welsch et al., 2010). To our knowledge, the *PSY2-Y-2* allele has been present in at least one copy in all yellow accessions and absent in all white accessions that have been genotyped at this locus. The *PSY2* locus is also negatively associated with dry matter content, thought to be driven by genetic linkage with an unfavorable allele for DM (Rabbi et al., 2017). There are two candidate genes for DM near *PSY2*: *sucrose synthase* and *UDP-glucose pyrophosphorylase* (Rabbi et al., 2017, 2020).

However, transgenic experiments have supported the hypothesis that *PSY2* has a pleiotropic effect on carbohydrate metabolism. Cassava and potato lines expressing a *PSY* transgene had 50-60% reductions in DM, increased sugar and fatty acids, and altered expression of carbohydrate metabolism genes compared to wild-type lines (Beyene et al., 2018). Similar shifts in primary metabolism have been observed in other crops engineered for higher carotenoid content, including tomato, maize, rice, and citrus (Li et al., 2012; Cao et al., 2015; Decourcelle et al., 2015; Gayen et al., 2016; Mi et al., 2022). Several hypotheses about a physiological impact of carotenoids on carbohydrate metabolism have been proposed; for example, the accumulation of phytoene may promote the conversion of amyloplasts (plastids that make and store both starch and TC) into chromoplasts for higher TC sequestration, reducing starch synthesis capacity (Ceballos et al., 2017; Beyene et al., 2018; Llorente et al., 2020; Villwock et al., 2024). A bivariate genome-wide association study (GWAS) for TC and DM in a west African yellow cassava breeding population showed that the genetic basis of the relationship between traits was polygenic, with many small-effect pleiotropic loci spread across the genome (Villwock et al., 2025). Understanding the physiological basis of this pleiotropy will inform breeding strategies for provitamin A biofortified starchy crops as well as provide insight into an interface of primary and secondary metabolism.

Previous studies have observed differential expression of key carotenoid biosynthesis/catabolism genes between white and yellow cassava varieties, including *1-deoxy-D-xylulose 5-phosphate synthase (DXS), lycopene ε-cyclase (LCYE)*,, *β-carotene hydroxylase 1 (BCH1)*, and *9-cis-epoxycarotenoid dioxygenase (NCED)* (Carvalho et al., 2016; Olayide et al., 2020, 2023; Wahyuni et al., 2020; Xiao et al., 2021). More recently, multi-omic studies have identified links between carotenoids and primary metabolism in cassava, although their findings differed. Among 49 African cassava genotypes, high carotenoid and low starch roots exhibited transcriptomic and metabolomic shifts towards myo-inositol, raffinose, and cell wall polysaccharide production (Gutschker et al., 2024). Proteomic and metabolomic analyses of eight African genotypes also associated high dry matter with a greater abundance of proteins involved in sugar conversion and glycolysis (Lamm et al., 2023). In contrast, a Latin American breeding population showed no significant negative correlation of carotenoids with dry matter or sugars, but *β*-carotene was positively correlated with plastid-associated lipids in the amyloplast fraction of eight selected genotypes (Drapal et al., 2024). Together, these studies point to relationships between carotenoids and primary metabolism, but do not distinguish genotype-specific effects or population-wide genetic correlations from general physiological interactions.

Some cassava varieties exhibit spatial variation in color within the storage roots, in which the inner tissue layer is often higher in TC than the outer layer (Chavez et al., 2008; Carvalho et al., 2016). The phenotype of a yellow inner tissue layer with a white or cream-colored outer tissue layer has been called “egg yolk” (Fig. 1). Spatial variation across tissue layers has been reported for DM as well (Chavez et al., 2008; Ceballos et al., 2012). Anatomically, these differences occur as concentric layers of storage parenchyma and vessels form during root secondary growth, with the outer tissue layer containing younger parenchyma closer to the cambium and the inner tissue layer containing older parenchyma and the vascular core (Chavez et al., 2008; Carvalho et al., 2016, 2017). Spatial variation in TC and DM across root tissues presumably reflects transcriptional, post-transcriptional, or other developmental regulation, since tissue layers share the same genotype. The population in this study contained roots with carotenoid enrichment in either the inner or outer layer, indicating that carotenoid accumulation and tissue layer were not completely confounded. The spatial variation therefore provided an opportunity to test for a physiological trade-off between TC and DM while accounting for genotype and tissue-layer effects. We reasoned that if the negative genetic correlation between TC and DM was driven by a physiological trade-off rather than (or in addition to) the genetic effects of the *PSY2* locus, then we would observe the negative correlation of TC and DM both across genotypes and across non-genetic (spatial, temporal, environmental) phenotypic variation.

**Figure 1.**
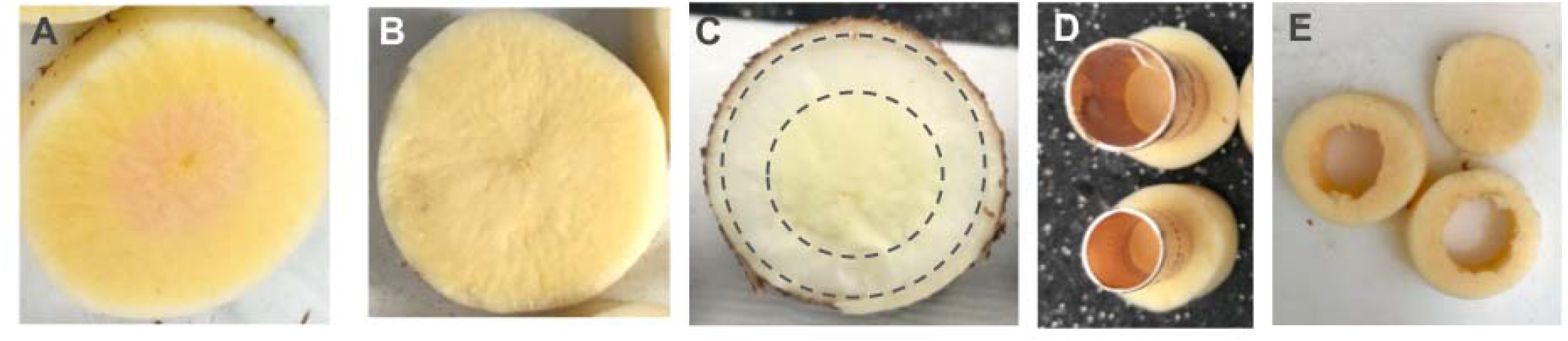
Images of peeled root cross sections with (A) or without (B) apparent spatial variation in visual color (a proxy for TC) between inner and outer tissue layers. C). Root cross section annotated with the ring where the approximated inner tissue layers were punched out with the rings shown in D). E). Outer tissue layer samples after punching out the inner tissue layer.

Partitioning phenotypic correlations into genetic and environmental components is common practice in multi-trait association and prediction. Likewise, considering gene expression as a phenotype, gene co-expression can be partitioned into genetic and non-genetic components. The genetic components of co-expression might be driven by, for example, a genetic variant in a transcription factor that upregulates a set of genes or in its binding site, effects that would be shared across clones of the same genotype. The non-genetic components of co-expression might be driven by a physiological response to an environmental condition or developmental differences between clonal replicates or roots of the same plant. To distinguish these effects, we extended linear modeling methods for estimating genetic and environmental effects to negative binomial regression models appropriate for RNAseq count data. We then integrated the resulting expression estimates with metabolite abundances in a joint gene-metabolite network using Weighted Gene Co-expression Network Analysis (WGCNA) (Langfelder and Horvath, 2008). While the WGCNA framework was originally developed for gene co-expression analysis, researchers have since applied it to metabolomic and proteomic datasets (Pei et al., 2017) and to combinations of -omics modalities (Umer et al., 2020; Wang et al., 2022). Joint gene-metabolite networks enable relationships between gene expression and metabolite variation to be evaluated within a common framework.

The main objectives of this study were to: 1) Investigate the transcriptional regulation of carotenoid accumulation and composition across genetic, temporal, and spatial variation; 2) characterize the network of interactions between carotenoids and carbohydrate metabolism; and 3) use this interaction network to evaluate support for mechanistic hypotheses to prioritize in future validation studies. By evaluating African and Latin American accessions in a common environment, quantifying carotenoid composition across root tissues and developmental stages, and partitioning genetic and non-genetic variation, this study distinguishes population-level genetic effects from inherent physiological relationships. Given the persistence of the genetic correlation and evidence of genome-wide pleiotropy between TC and DM (Villwock et al., 2025), we hypothesized that carotenoid and starch metabolism genes would be strongly connected in both genetic and non-genetic co-expression networks, with abscisic acid signaling and/or plastid development genes as potential connectors.

## Materials and Methods

### Plant material and sample collection

The experimental design is summarized in Fig. 2. We started with a population of 5 African cassava (*Manihot esculenta* Crantz) accessions, 5 Latin American accessions, and 44 African x Latin American progeny from crosses between them made at the NextGen Cassava Project crossing nursery at the University of Hawai’i at Mānoa. One parent from each population was a white-fleshed accession, while the remainder exhibited a range of yellow root colors from light cream to intense yellow. The hybrid progeny were grown from botanical seed and subsequently clonally propagated alongside stem cuttings of the parental accessions in 2022 (year 1). Three replicates of each parent and un-replicated progeny were planted in 25cm pots in an augmented randomized block design with the parents as replicated checks at the Guterman Bioclimatic Laboratory greenhouse in Ithaca, NY under temperature, light, and media conditions as in Hyde et al. (2020). Plants were fertilized biweekly with a balanced fertilizer until 4 months after planting (MAP), when the fertilizer was switched to a high phosphorous fertilizer, Jack’s Blossom Booster 10-30-20 (JR Peters Inc.), to promote storage root formation as in Taylor et al. (2012). At 4 MAP, plants were pruned back to 1.3m height; any plants shorter than this height at the time of pruning had the top apical meristem snipped off.

**Figure 2.**
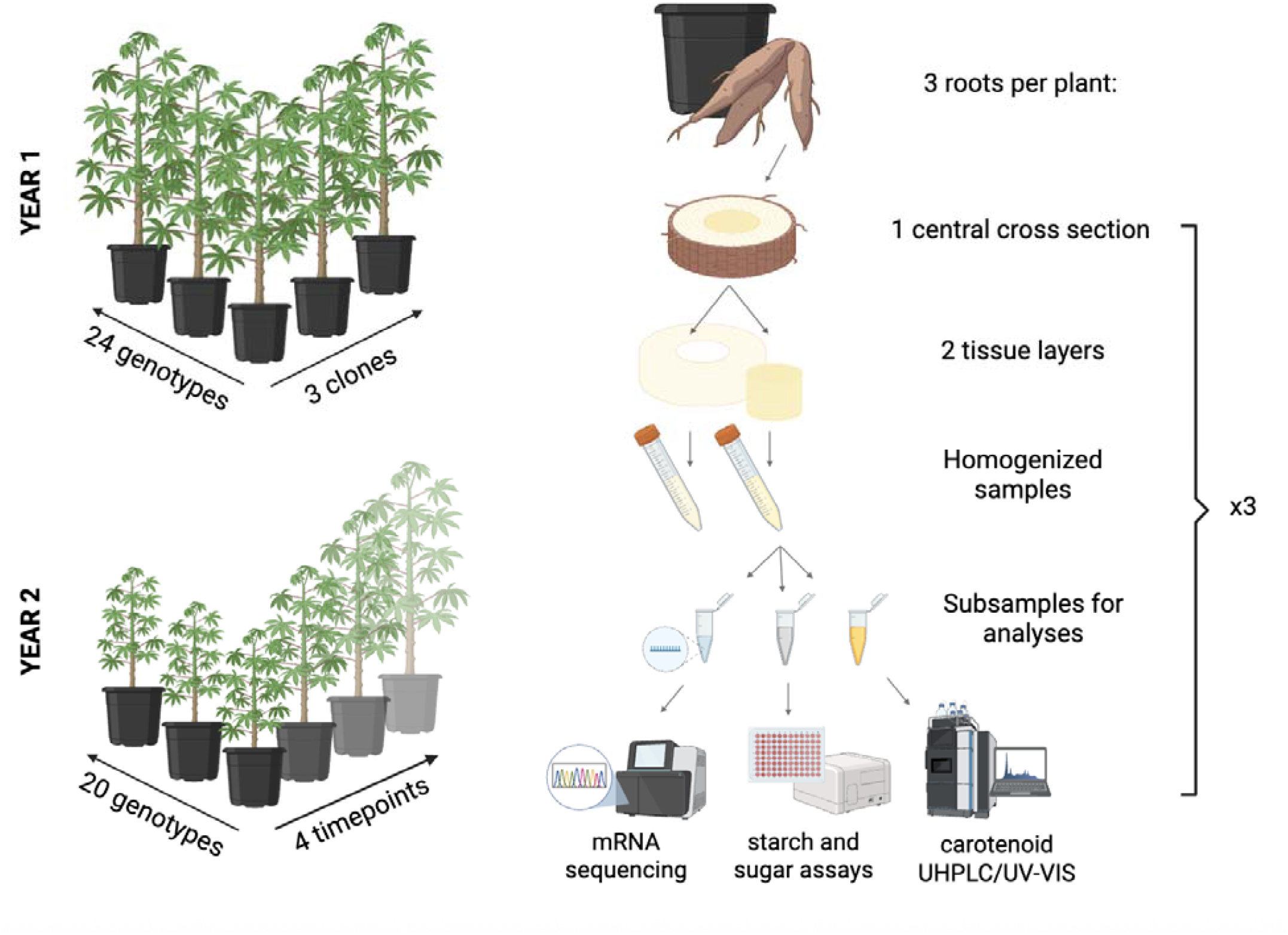
Overview of the experimental design, sampling, and analysis methods.

Harvesting was conducted at 5 MAP by pulling out the three largest storage roots per pot. Each root was rinsed and peeled, and a single cross section about 2cm thick was cut out at the midpoint of the root length. The inner and outer root tissue layers were separated using copper pipe couplers to punch out the layer of the peeled root cross section as in (Carvalho et al., 2017). The diameter of the coupler (ranging from 0.6 to 2.5cm) was selected to most closely fit the visible boundary of storage parenchyma pigmentation when apparent, or approximated proportional to root size as shown in FigureD. Therefore, the inner and outer tissue layer samples represent approximate spatial zones within the storage parenchyma rather than precise tissue dissections. Root tissue samples were quickly chopped, placed into a 15mL tube, weighed, flash frozen in liquid nitrogen, and kept on dry ice until placed in a -80° C freezer.

A subset of 20 genotypes representative of spatial variation in carotenoid content were re-planted in 2023 (year 2) for replicated evaluation across a series of harvest timepoints (Table S1). Stem cuttings were planted in a randomized complete block design across 4 greenhouse blocks and grown as described above, with pruning monthly beginning at 4 MAP. At each harvest timepoint (6, 7, 8, and 9 MAP), one plant replicate of each genotype was selected randomly with respect to blocks. The remaining un-harvested plants were pruned at each harvest timepoint such that harvesting was always conducted one month after last pruning.

Frozen root tissue samples were lyophilized for 48 hours and dry weight recorded, then placed back into the -80°C freezer. Dried, frozen root samples were ground and homogenized using pre-chilled steel grinding beads in a Spex SamplePrep Geno/Grinder at 1400 rpm for four 15-second intervals, placing the tubes back in liquid nitrogen for at least 1 minute between grinding intervals to keep the tissue frozen. Ground samples were placed back in the -80° C freezer until further processing.

### Carotenoid extraction and quantification

Each carotenoid extraction batch included two replicate extractions from a standardized sample to assess technical variation across batches. To prevent light- and heat-induced degradation or isomerization of carotenoids, sample racks were wrapped in foil and kept on ice throughout the procedure. Carotenoids were extracted using a modified protocol from Li et al. (2012). Briefly, 1 mL of cold ethyl acetate was added to ∼250 mg of ground root tissue, which was then vortexed with two 2 mm steel grinding beads at 4°C for 30 minutes. After centrifugation, the supernatant was collected, and the extraction was repeated three additional times on the same tissue. The resulting extracts were pooled and back-extracted by washing with 3 mL of 0.1 M NaCl, followed by vortexing for 3 minutes at 4°C. The upper organic phase was transferred to a clean tube, dried under vacuum centrifugation, and stored at - 20°C until Ultra High Performance Liquid Chromatography (UHPLC) analysis.

For UHPLC analysis, the dried samples were reconstituted in ethyl acetate containing 0.05 mg/mL diindolylmethane (DIM), used as an internal standard. Carotenoids were separated and quantified using ultra-performance convergence chromatography (UPC^2^), adapted from (Gonda et al., 2019). Chromatographic separation was performed on a Waters HSS C18 SB column (3.0 × 150 mm, 1.8 µm particle size) at 1.0 mL/min and 40°C. Detection was carried out using a photodiode array detector, monitoring wavelengths from 250 to 700 nm.

The column was equilibrated with 99% eluent A (supercritical CO□) and 1% eluent B (methanol). Carotenoids were eluted using a nonlinear, concave gradient that ramped to 20% B over 7.5 minutes, held for 4.5 minutes, then returned to 1% B, followed by 3 minutes of re-equilibration. Individual carotenoids were identified by comparison to authentic standards (Sigma) and quantified using MassLynx software v4.1 (Waters) at compound-specific wavelengths chosen for optimal sensitivity and selectivity. A five-point β-carotene calibration curve (5–100 ng/mL) was used for quantification, and all concentrations were reported as β-carotene equivalents. Chromatographic details are presented in Fig. S1.

### Starch and sugar quantification

Dry matter content was calculated as the ratio of dry weight to fresh weight measurements (before and after lyophilization) after subtracting out the average weight of an empty 15mL tube. Carbohydrate composition was additionally analyzed in the samples from year 1 only. Glucose, sucrose, and starch were quantified by peroxidase glucose oxidase (PGO) colorimetric assay (Trinder, 1969; Lott and Turner, 1975) protocol adapted from (Setter et al., 2001). Sugars were extracted from 20mg of powdered root sample with 1mL 80% ethanol at 60°C for one hour and centrifuged. Two replicates of each supernatant were reacted with PGO reagent (1 U/mL peroxidase, 2 U/mL glucose oxidase, 9 mg/mL para-hydroxybenzoic acid, 0.3mg/mL 4-aminoantipyrene, pH 4.5) and incubated one hour at room temperature. Absorbance at 500nm was measured using an Epoch 2 microplate spectrophotometer (BioTek Instruments, Inc.). Glucose concentrations were calculated by linear regression from glucose standards on each plate (Fig. S2). Outliers were removed if Cook’s distance was more than four times the mean. For sucrose, extracts were first incubated with invertase (0.75mg/mL) for two hours at room temperature. Glucose equivalents were then measured with PGO and sucrose content was calculated as (total - free glucose) x 1.9. The remaining pelleted tissue was washed with another 80% ethanol extraction, gelatinized in water at 100°C for 4 hours, then digested with amyloglucosidase (105 U/mL) and α-amylase (5 U/mL) in acetate buffer for 24 hours at 45°C. After confirming complete starch hydrolysis with Lugol’s iodine solution, digests were diluted and assayed with PGO. Starch content was calculated as glucose equivalents x 0.9 x dilution factor.

### Phenotypic data analysis

Outliers were identified and removed using *outliers*∷*grubbs*.*test()* in R. Kendall’s tau correlation method was used to test correlations between phenotypes since they exhibited a right-skewed distribution. Phenotypic associations between carotenoids and dry matter were then tested with linear mixed models fit with maximum likelihood using *lme4* in R:

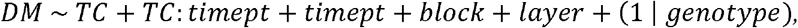

where *DM* is the ratio of dry weight to fresh weight, *TC* is total carotenoids in μg/g dry weight, *timept* is the fixed effect of the harvest timepoint factor, *TC:timept* is the fixed effect of the interaction between TC and harvest timepoint, *block* is the fixed effect of the greenhouse bench, *layer* is the fixed effect of inner vs. outer root tissue layer, and (1 |*genotype*) is the random effect of genotype. The effect of TC and the interaction effect between TC and timepoint on DM were tested for statistical significance by likelihood ratio comparison between the full model and a reduced model without the *TC* and *TC:timept* covariates.

To examine the relative contributions of different carotenoids to the association with dry matter, carotenoids were partitioned into total *cis-*carotenes (the linear upstream carotenes: phytoene + phytofluene + ζ-carotene) and total other carotenoids (the sum of all remaining carotenoid metabolites: lutein + β-carotene + violaxanthins) as separate covariates in the following linear model:

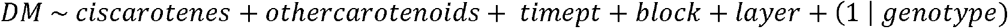

The contribution of *cis-*carotenes was tested by likelihood ratio comparison between nested mixed models with and without the *cis-*carotenes covariate. The effects of individual carotenoid metabolites on DM were also tested in the same manner, replacing the *cis-*carotene term with the individual carotenoid and calculating “other carotenoids” as the sum of all remaining carotenoids.

To test whether the relationship between TC and DM differed between African and Latin American cassava accessions, the following linear model was fit:

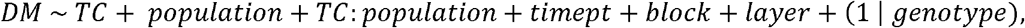

where *population* is a categorical variable indicating African, Latin American, or hybrid origin. The interaction of TC and population origin was tested by likelihood ratio comparison between nested mixed models with and without the *TC:population* interaction term.

### RNA extraction and sequencing

Subsets of 190 samples were selected from each year for transcriptome profiling. From year 1, all tissue layer samples from the 20 genotypes selected for advancement and the remaining 5 parents were selected. From year 2, all samples from replicates harvested at 9 MAP were selected for transcriptome profiling, since this timepoint exhibited the strongest negative correlation between TC and DM. Additional samples from 4 representative genotypes across all timepoints were selected to capture temporal variation. The genotypes C17, X1_54, X11_68, and X2_24 were selected as temporal representatives based on capturing the overall phenotypic trends across timepoints, as assessed by the lowest total distance between the genotype/layer points and linear model predictions for TC and DM across timepoints and genotypes. Therefore, the subset for transcriptomic analysis contained 25 genotypes, summarized in Table S1.

Total RNA was extracted from the selected sample subset using the hot borate method (Wan and Wilkins, 1994) modified as described in Hershberger et al. (2022). RNA samples were treated with the Invitrogen TURBO DNA-free Kit (Thermo Fisher Scientific) to remove residual genomic DNA. RNA yield, quality, and integrity were evaluated by agarose gel electrophoresis and using an Epoch 2 microplate spectrophotometer (BioTek Instruments, Inc.). Samples with less than 100 ng/μL concentration or less than 1.9 A260/280 ratio were re-extracted or excluded. Samples passing quality thresholds were diluted to a concentration of 100-500 ng/μL and plated in 96-well plates (2 plates in 2022 and 2 in 2024).

RNA sequencing was conducted by the Biotechnology Resource Center Genomics Facility at the Cornell Institute of Biotechnology (RRID:SCR_021727). Library preps were done with the Lexogen QuantSeq V1/V2 kits. Year 1 samples were sequenced with NextSeq 500/550 single-end (1x75bp) and year 2 samples were sequenced with NovaSeqX paired-end (2x150bp), though only read 1 was used for downstream analyses. Fastq files were generated by the Illumina pipeline software v2.18.

### RNAseq data analysis

Year 1 RNAseq data (NextSeq) consisted of an average of 4.7 million single-end reads per sample, while year 2 data (NovaSeqX) had an average of 8.2 million paired-end reads per sample. Read quality was checked with FastQC/MultiQC (Andrews, 2010; Ewels et al., 2016). NextSeq (year 1) reads were trimmed with cutadapt (Martin, 2011) to remove Illumina adapters, poly-A and poly-G sequences of at least 10 bases (characteristic of “no signal” on the 2-channel NextSeq instruments), low-quality bases (<10 phred score) from the 3’ end, and the first 12 bp on the 5’ end (corresponding to the random primer), keeping only reads at least 18 bases in length. NovaSeqX (year 2) reads were trimmed with cutadapt using the same parameters except for the poly-G trimming specific to NextSeq instruments.

Trimmed reads were aligned to *M. esculenta* reference genome v8.1 (Bredeson et al., n.d.; Goodstein et al., 2012) and counted using STAR aligner (Dobin et al., 2013) with the following parameters: maximum number of multiple alignments allowed per read = 10, maximum number of mismatches = 10, remove non-canonical unannotated splice junctions, maximum intron size = 20,000, maximum ratio of mismatches to mapped read length = 0.06. After alignment, samples with poor mapping (< 100,000 uniquely mapped reads) or overall low expression (< 10,000 genes expressed with 10 or more counts) were filtered out, leaving 306 samples passing the quality thresholds (77% of year 1 samples and 84% of year 2 samples). Genes that were expressed (nonzero counts) in less than 20 samples in each of the NextSeq and NovaSeqX datasets were filtered out, leaving 20,430 expressed genes present in both datasets that were then combined into a single matrix. Data structure was visualized with *plotPCA()* (Fig. S3) after variance-stabilizing transformation of raw gene counts with *vst()* (Fig. S4) using the DESeq2 R package (Love et al., 2014).

Given the different sequencing methods across years, there was substantial stratification of the raw RNAseq data by batch (Fig. S3). To remove batch effects and other sources of technical variation that could confound co-expression patterns, surrogate variable analysis was conducted using the *SVA* package in R (Leek et al., 2012). Surrogate variable analysis (SVA) uses an algorithm to estimate latent factors associated with broad variation across all genes while preserving the effects of variables of interest (Leek and Storey, 2007). In this case, we assume that the major components of error variation collectively across all genes capture technical factors rather than biological variation, which would be more likely to affect only subsets of certain interacting genes. Briefly, the SVA algorithm fits a linear model with known covariates of interest and conducts a singular value decomposition on the residual matrix to extract residual eigengenes, which are then tested for significance by comparing their variance explained to a null distribution derived from a permuted residuals matrix that removes structure across samples. SVA was performed with the following generalized linear model and a corresponding reduced model with only an intercept term using the *svaseq()* function, which uses a log-link to accommodate gene count data:

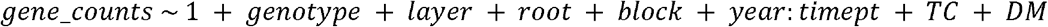

Known sources of biological variation and the key phenotypes of interest (TC and DM) were included as covariates in the model to prevent SVA from capturing biological factors of interest. Missing phenotype data for total carotenoids (4 samples) and dry matter content (6 samples) were imputed using the mean values for the given genotype and tissue layer. To determine how many of the surrogate variables to include as covariates in downstream analyses, the significant surrogate variables were included as covariates in a negative binomial model fit with DESeq2 (Love et al., 2014) and the resulting p-value distributions were examined. The top seven surrogate variables were selected as covariates since they had p-value distributions strongly skewed towards zero, representing significant associations with most genes (Fig. S5).

### Characterizing carotenoid trait-associated genes

Trait-associated genes (TAGs) significantly associated with overall phenotypic variation in TC, after accounting for differences between tissue layers and the effects of surrogate variables on transcript abundance, were identified with the following negative binomial model design in DESeq2, where SV represents the surrogate variables:

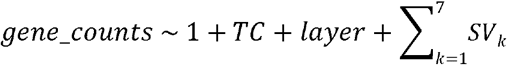

TAGs were further partitioned into genetic and non-genetic TAGs using separate models that tested the effect of TC while controlling for different sources of transcript variation. Genetic TAGs, genes associated with TC after accounting for the effects of non-genetic covariates on transcript abundance, were identified with the following DESeq2 design:

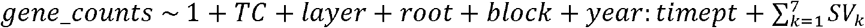

Non-genetic TAGs, genes associated with TC after accounting for the effects of genotype and tissue layer on transcript abundance, were identified with the following DESeq2 design:

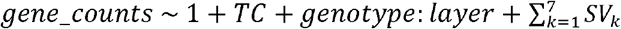

Egg-yolk TAGs, genes associated with the egg-yolk phenotype (measured continuously as the difference in total carotenoids between the inner and outer tissue layer) and the interaction effect of egg-yolk and inner tissue layer, were identified with the following DESeq2 design:

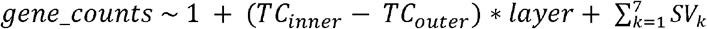

P-values were adjusted with the Benjamini & Hochberg (1995) false discovery rate adjustment for multiple test correction. TAGs were classified as statistically significant if the effect estimate of the TC trait was nonzero with an adjusted p-value < 0.05.

### Ontology enrichment

Sets of TAGs with |log_2_-fold change| (LFC) values greater than 0.5 were tested for enrichment of GO terms using the *R* package *clusterProfiler* (Wu et al., 2021). Ontology term enrichment was conducted with *enricher()* using custom GO annotations from the *M. esculenta* v8.1 assembly obtained from Phytozome (Bredeson et al., n.d.; Goodstein et al., 2012). All expressed genes tested for differential expression were used as the background gene set. P-values were adjusted for multiple testing using the Benjamini-Hochberg (1995) correction. Ontology terms were considered significantly enriched if the adjusted p-value of the hypergeometric test was less than 0.05. Ontology term enrichment was visualized with *enrichplot::dotplot*.

### Partitioning gene expression and metabolite phenotypes into genetic and non-genetic components

Variation in gene expression was partitioned into genetic and non-genetic components using the following negative binomial generalized linear model fit with DESeq2:

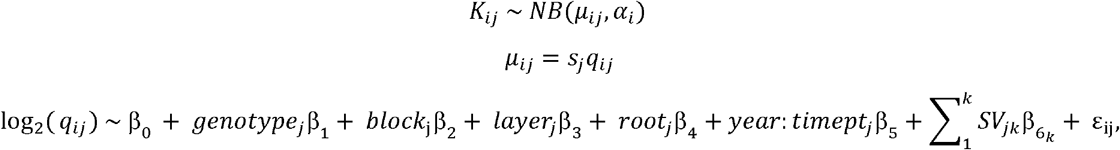

where *k*_*ij*_ represents the observed counts of gene *i* for sample *j* modeled with a negative binomial distribution with mean *µ*_*ij*_ and gene-specific dispersion parameter *α*_*i*_, *s*_*j*_ is the size normalization factor for sample *j, q*_*ij*_ is the estimated true expression proportion for gene *i* in sample *j*, and *q*_*ij*_ is modeled on the log_2_ scale with a linear model with intercept β_0_, fixed effect estimates β_1-5_ for experimental variables (genotype, greenhouse block, tissue layer, root number, and harvest timepoint within year), the sum of *k* surrogate variables *SV*_*j*_ and their associated coefficients 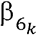, and residual error ε_ij_ for gene *i* in sample *j*.

The vector of log_2_ -scale residuals ε was calculated as follows:

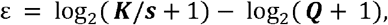

where ***K/s*** is the matrix of size-normalized observed gene counts and ***Q*** is the matrix of fitted values *q*_*ij*_, with a pseudocount of 1 added to handle genes with zero counts.

The best linear unbiased estimates (BLUEs) of overall expression, the genetic effects on expression, and the non-genetic effects on expression (normalized on a log_2_-like scale) were then calculated by summing the relevant coefficients extracted from the model as follows:

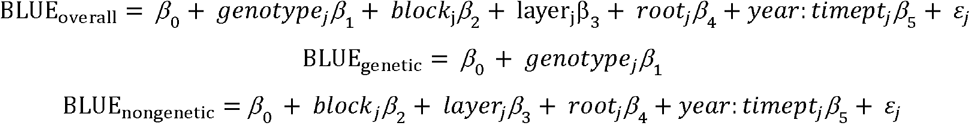

The outputs were three matrices of batch-corrected gene expression estimates: overall (*n*_*samples*_ *x g*), genetic (*n*_*genotypes*_ *x g*), and non-genetic (*n*_*samples*_ *x g*), where *g =* 20,430 genes, *n*_*samples*_ = 306 samples, and *n*_*genotypes*_ = 24 genotypes.

Likewise, for the carotenoid, dry matter, and carbohydrate data, the metabolite estimates were log_2_ transformed to match the scale of the gene expression estimates as described in Pei et al., (2017). The batch-corrected genetic and non-genetic estimates of metabolite abundance were then extracted as described above from the following linear model fit with *lm()* in R:

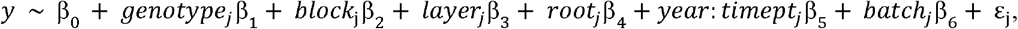

where *batch* is the batch processing date for carotenoid extractions, lyophilization, or carbohydrate analysis for the relevant metabolite *y* on a log_2_-scale for sample *j*, and other variables are as defined above. These log_2_-scale genetic, non-genetic, and overall (batch-corrected) estimates of metabolite abundance were then added as columns in the matrices of the corresponding gene expression estimates.

### Gene-metabolite network construction

The three gene/metabolite matrices of overall phenotypic, genetic, and non-genetic effect estimates were then input separately for network construction with the WGCNA (Weighted Gene Co-expression Network Analysis) package in R (Langfelder and Horvath, 2008). Outliers were checked with the *goodSamplesGenes()* function. The key parameters used in WGCNA were unsigned network type (treating both positive and negative correlations as evidence of connectivity) and the use of biweight midcorrelation (bicor) instead of Pearson correlation to improve robustness to outliers. The *pickSoftThreshold()* function was used to select the lowest power β for the adjacency calculation to approximate a scale-free topology model fit (R^2^ ≥ 0.85): 7 for the overall network: 10 for the non-genetic network, and 6 for the genetic network (Fig. S6). Key parameters used in network construction were a minimum module size of 30, deepSplit of 3, module reassignment threshold of 0.05, and merge cut height of 0.25, to balance between module number and size.

The resulting matrices of topological overlap measures (TOMs) represented the genetic, non-genetic, and overall connection between each pair of genes or metabolites as a function of adjacency and shared neighbors:

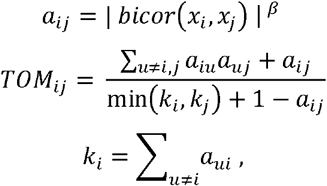

where *a*_*ij*_ is the pairwise adjacency between estimate *x* of genes and/or metabolites *i* and *j, β* is the soft thresholding power, *TOM*_*ij*_ is the topological overlap measure between genes and/or metabolites *i* and *j*, and *k*_*i*_ is the connectivity of gene/metabolite *i* with all other genes/metabolites *u* (Langfelder and Horvath, 2008). The resulting TOM matrices calculated from the genetic, non-genetic, and overall estimates of gene/metabolite abundances will hereafter be referred to as TOM_G,_ TOM_E,_ and TOM_G+E_, respectively.

### Carotenoid and starch metabolism interactions

Gene annotations for the *M. esculenta* reference genome v8.1 were obtained from Phytozome (Bredeson et al., n.d.; Goodstein et al., 2012). Genes were classified *a priori* as related to carotenoid metabolism if they had MetaCyc pathway annotations matching carotenoid and apocarotenoid biosynthesis, carotenoid cleavage, geranylgeranyl diphosphate biosynthesis via the methylerythritol phosphate (MEP) pathway, or abscisic acid biosynthesis, or gene ontology (GO) terms for carotenoid metabolic process, carotenoid biosynthetic process, isoprenoid metabolic process, regulation of isoprenoid metabolic process, chromoplast, or any subclasses within (63 genes; Table S2). Genes were classified as related to starch metabolism if they had MetaCyc pathway annotations matching the superpathway of sucrose and starch metabolism I (non-photosynthetic tissue), polysaccharide biosynthesis, starch biosynthesis, starch degradation, sucrose degradation, glycolysis, or pyruvate dehydrogenase and TCA cycle, or GO term annotations matching starch metabolic process or its regulation, glycolytic process or its regulation, maltose metabolic process, hexose transmembrane transporter activity, response to hexose, carbohydrate storage, amyloplast, starch grain, or any subclasses within (240 genes; Table S2).

To assess whether these gene sets were more connected in the co-expression networks than would be expected by chance, the total TOM between all pairs of genes in the carotenoid and carbohydrate gene sets was compared to an empirical null distribution generated via node permutation (McCormack et al., 2013). Metabolite nodes were excluded from this analysis, since only biologically relevant metabolites were measured. To generate an appropriate null distribution for total TOM, a set of random genes were selected that had approximately the same overall connectivity *k* as genes in the carotenoid-related gene set. Gene connectivities were binned into 10 quantiles; for each carotenoid gene, a random gene not in the either gene set was drawn from the corresponding connectivity quantile, such that the random gene set reflected the size and overall connectivity of the carotenoid gene set. The total TOM between the random gene set and the carbohydrate gene set was then calculated by subsetting the full TOM matrix, and the permutation was repeated 100,000 times. An empirical p-value was calculated as the proportion of times the observed TOM sum between carotenoid and carbohydrate gene sets was exceeded by that of random genes with the carbohydrate gene set. The permutation test was then repeated but with the carotenoid genes fixed and random gene sets selected to match the overall connectivity of the carbohydrate gene set. Permutations were run separately with the TOM_G+E_, TOM_G,_ and TOM_E_ matrices.

The strongest connections between carotenoid and carbohydrate metabolism were identified as those with the highest pairwise TOM. Modules most associated with carotenoid and carbohydrate metabolites from each of the three networks were visualized with Cytoscape (Shannon et al., 2003). The intramodular hub genes from each of these modules were identified as those with the highest correlation with the module eigengene (Langfelder and Horvath, 2008). The genetic and non-genetic relationships with carotenoid and starch genes/metabolites were compared by using a Mantel test (Mantel, 1967) of matrix correlations on the dissimilarity matrices (1-*TOM*_*G*_ and 1-*TOM*_*E*_), implemented with the *ecodist* R package with 1,000 permutations (Goslee and Urban, 2007).

## Results

### The carotenoid-dry matter relationship depends on developmental stage and root tissue layer

In the year 1 experimental trial, with roots harvested at 5 MAP, the overall relationship between TC and DM was slightly positive (τ = 0.105, p = 0.001) across tissue layers. However, the direction of the correlation differed between the inner and outer tissue layers, with the inner layer showing a non-significant negative correlation (τ = **-**0.088, p = 0.053) and the outer layer showing a significantly positive correlation (τ = 0.249, p = 4.5×10^-8^) (Fig. 3A). In the year 2 experimental trial, harvested at 4 timepoints 6-9 MAP, the overall relationship between TC and DM across timepoints was significantly negative (τ = -0.155, p = 6.9×10^-7^), with a stronger negative relationship in the inner tissue layer (τ = -0.222, p = 4.5×10^-7^) and an insignificant relationship in the outer layer (τ = -0.070, p = 0.115) (Fig. 3B). The relationship between TC and DM became increasingly negative across timepoints, from τ = -0.006 at 6 MAP to τ = -0.407 at 9 MAP (Fig. 4). There was a significant interaction effect between TC and timepoint on DM after accounting for genotype, block, and layer in a mixed linear model (*x*^2^(3)=14.9, p = 0.002). The estimated effect size of TC on DM indicated that each 1 μg/g TC corresponded to a - 0.05% decrease in DM on average across months.

**Figure 3.**
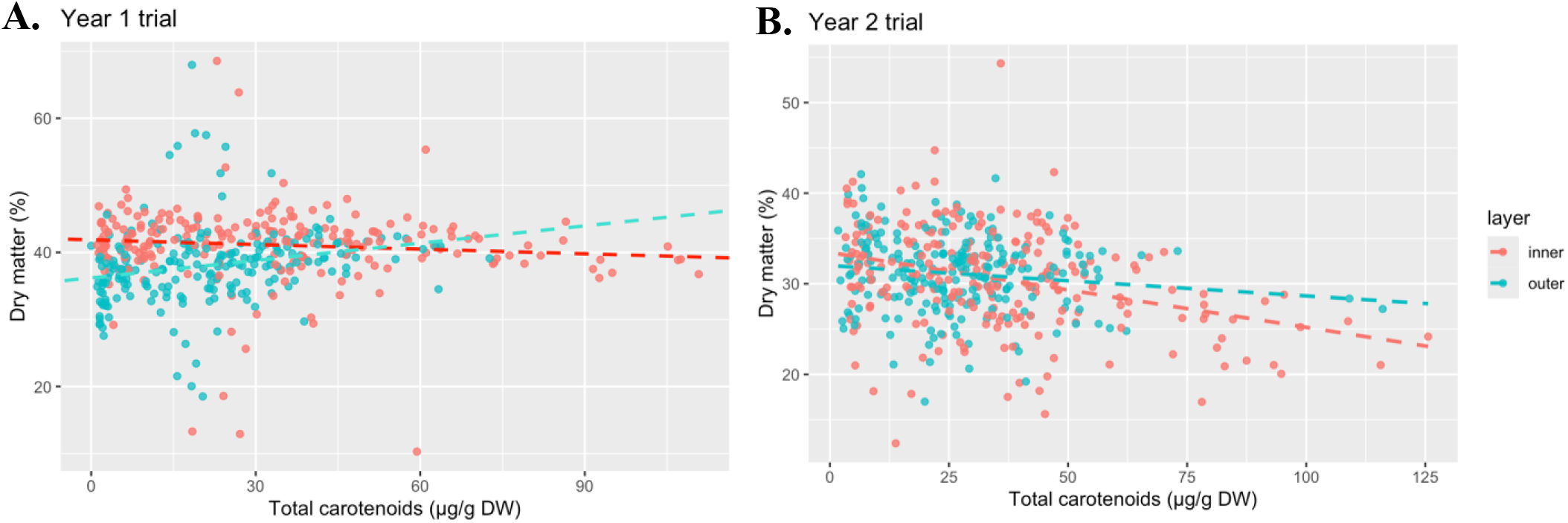
Phenotypic relationship between total carotenoids and dry matter content among the inner root tissue layer samples (red) versus outer tissue layer (blue) in the year 1 (left) and year 2 (right) trials. Each point represents an individual root tissue layer sample colored by tissue layer. Dashed lines represent the linear regressions of DM on TC for each tissue layer.

**Figure 4.**
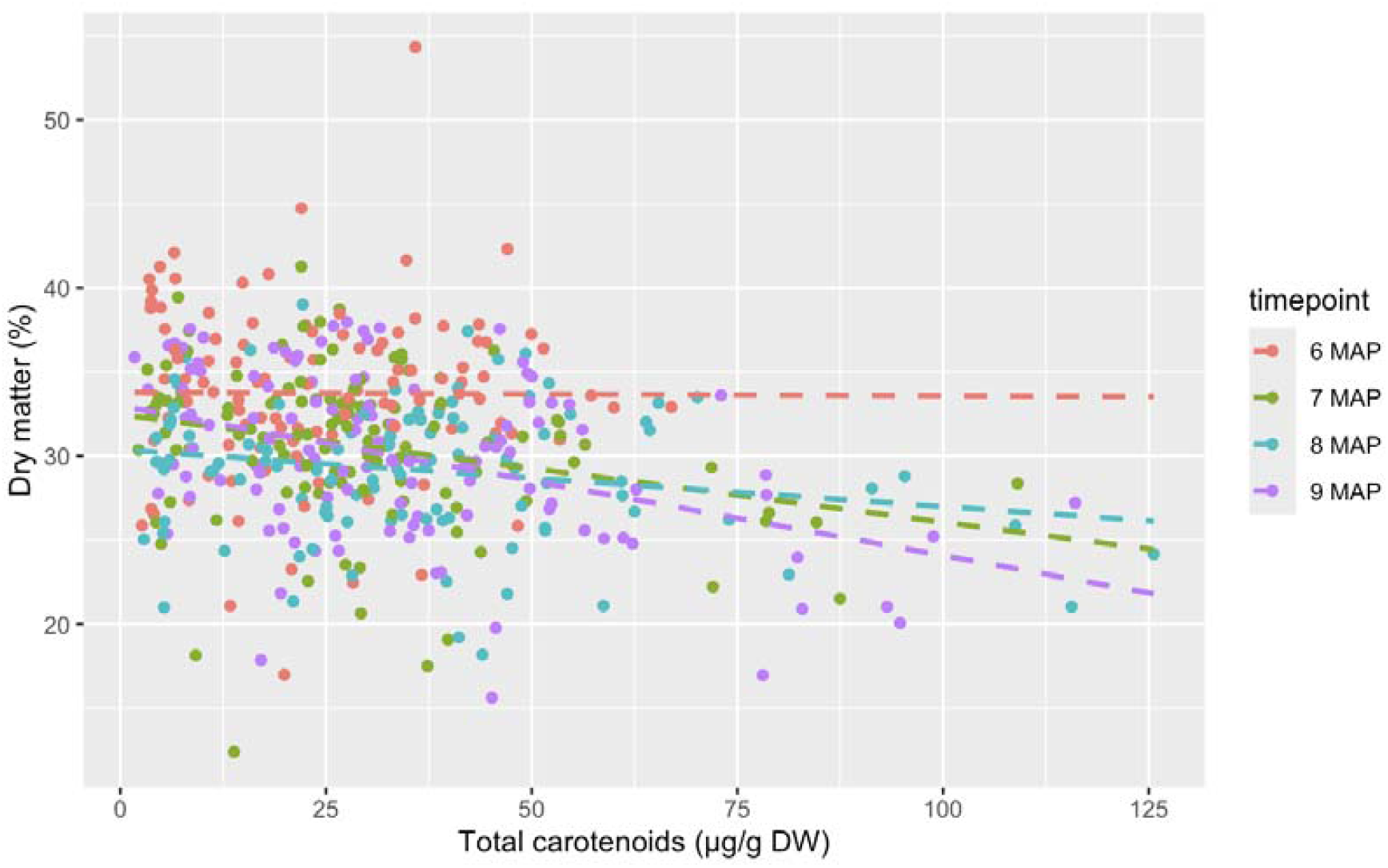
The relationship between TC and DM became increasingly negative across harvest timepoints in the year 2 trial. Each point represents an individual root tissue layer sample colored by the harvest timepoint. Dashed lines represent the linear regressions of DM on TC for each timepoint.

Across years, the relationship between TC and DM differed slightly among samples from African and Latin American accessions. In the mixed linear model accounting for genotype, year/timepoint, block, and tissue layer, the effect of TC on DM was less negative in Latin American samples than in African samples, with the estimated TC slope differing by 0.08. Including the interaction effect between TC and population of origin significantly improved model fit (*x*^2^(2)=9.24, p = 0.010). However, there were only five African and five Latin American parental accessions, which are not necessarily representative of the overall germplasm pools.

### Glucose content is negatively correlated with both dry matter and carotenoids

Of the correlations between carbohydrates and carotenoids (Fig. S8), the most prominent relationships were negative correlations between both glucose and DM (r = -0.51, p = 3×10^-21^; Fig. 5A) and glucose and TC (r = -0.23, p = 7×10^-5^; Fig. 5B). Starch content expressed on a fresh weight basis (mg/g dry weight * DM/100) was positively correlated with DM percentage (r = 0.59, p = 9×10^-42^; Fig. S7), as expected given that cassava DM is known to be comprised mainly of starch. Correlations between starch and TC in the inner and outer tissue layers followed the same directions of the DM and TC relationships respectively but were weaker, with a positive correlation in the outer tissue layer and no correlation in the inner tissue layer.

**Figure 5.**
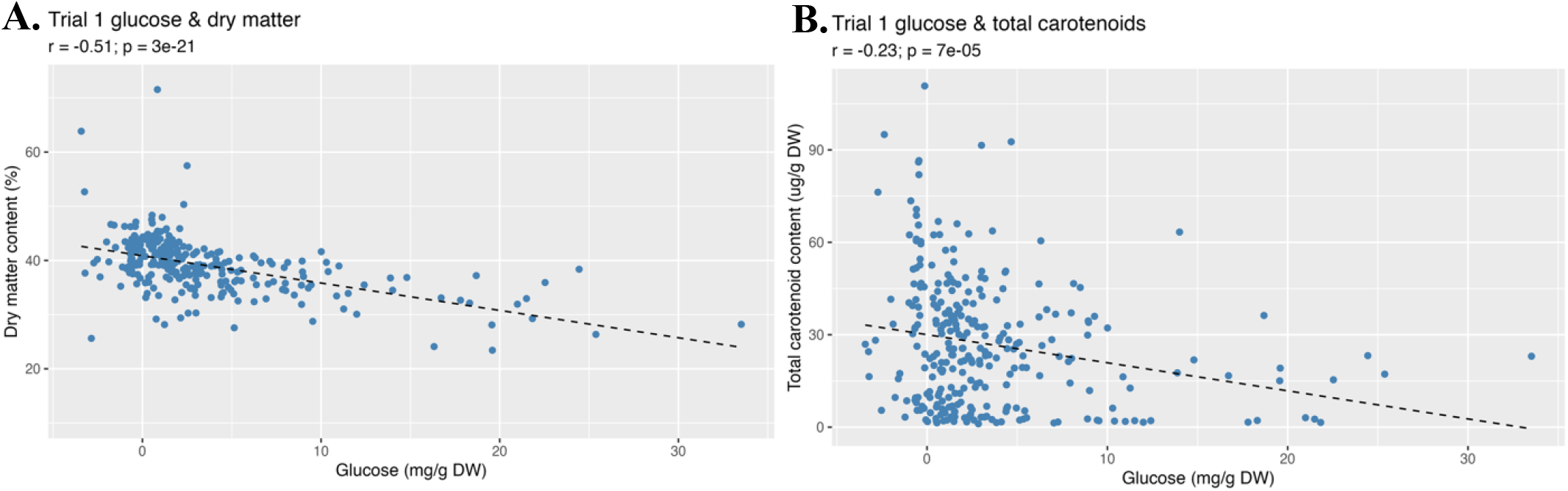
Correlations between glucose and DM (left) and glucose and TC (right). Each point represents a mean value per sample across two technical replicates. Dashed lines represent the linear regressions of DM on glucose and TC on glucose, respectively.

### Upstream cis-carotenes, a minor component of TC, are most strongly associated with dry matter

The identifiable carotenoid compounds were phytoene, phytofluene, *9-cis-*β-carotene and *all-trans*-β-carotene (analyzed jointly as total β-carotene), lutein, and two violaxanthin isomers (analyzed jointly as total violaxanthin). β-carotene was the primary carotenoid, comprising 84% of TC on average, with 8% upstream *cis-*carotenes (phytoene, phytofluene, and ζ-carotene), 6% violaxanthin, and 2% lutein. Principal component analysis separated carotenoid composition primarily by β-carotene along the first axis and by upstream *cis-*carotenes versus downstream xanthophylls (lutein and violaxanthin) along the second axis (Fig. 6).

**Figure 6.**
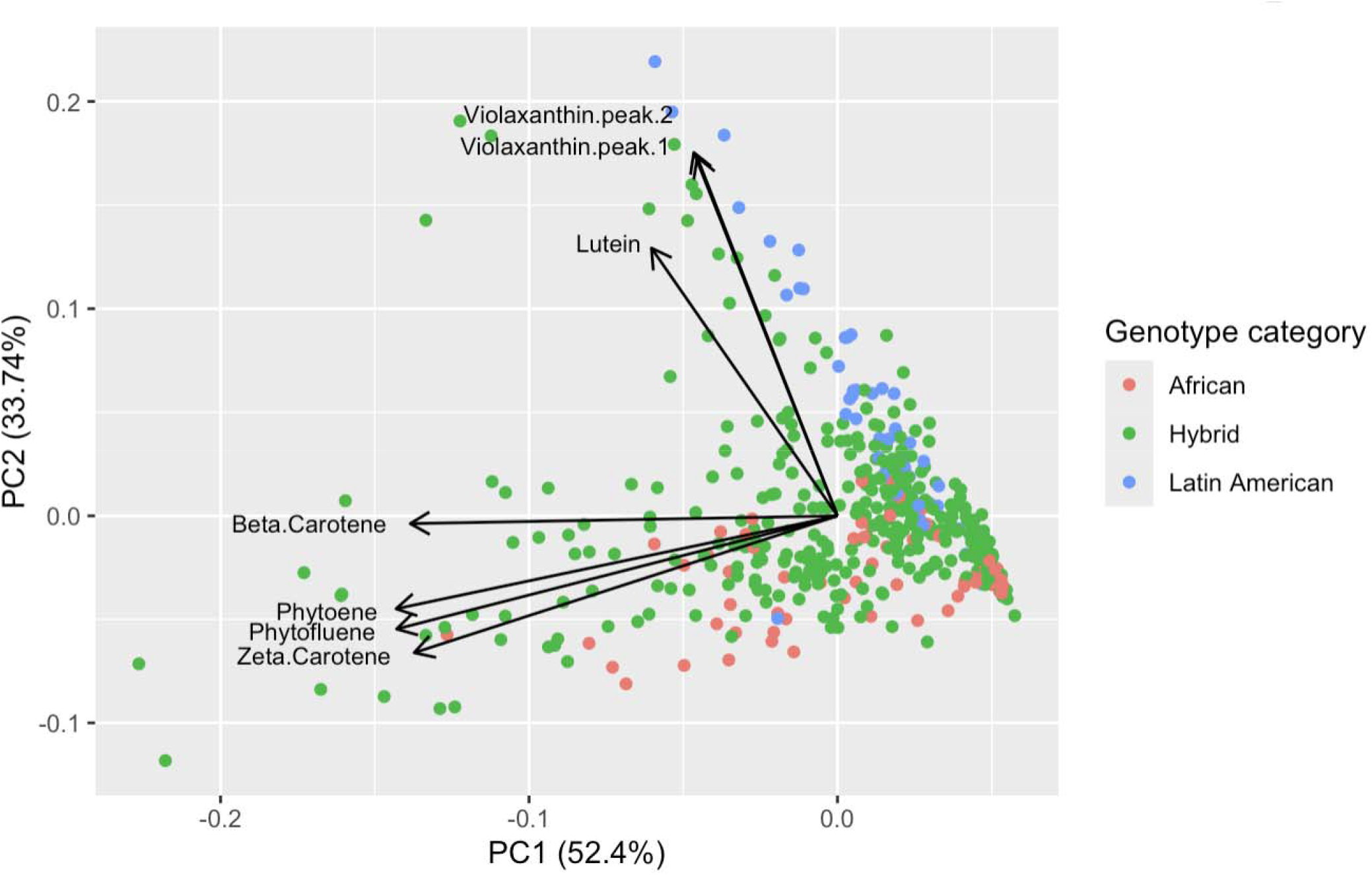
Principal component analysis of carotenoid composition. Points represent individual root tissue-layer samples and are colored by population of origin. Arrows show the loadings of carotenoid compounds on the first two principal components.

In a mixed linear model partitioning carotenoid composition into total *cis-*carotenes and total other carotenoids (β-carotene, lutein, and violaxanthins), *cis-*carotenes were negatively associated with dry matter after accounting for genotype, timepoint, tissue layer, and block (β = -0.34, p = 0.00014), while the effect of total other carotenoids was not significant (β = 0.03, p = 0.14). Adding total *cis-* carotenes significantly improved model fit relative to a reduced model containing total other carotenoids but not *cis-*carotenes (likelihood-ratio test, *x*^2^(1)= 12.8, p = 0.00034), indicating that upstream *cis-*carotenes explained additional variation in DM beyond what was captured by the other carotenoid metabolites. Similarly, adding phytoene or phytofluene individually significantly improved model fit relative to reduced models containing all other carotenoids (phytoene: *x*^2^(1) = 17.0, p = 3.68 × 10^-5^; phytofluene: *x*^2^(1) = 10.2, p = 0.0014), but ζ-carotene, β-carotene, lutein, and violaxanthin did not (p > 0.05).

### Carotenoid accumulation is most strongly associated with regulatory genes, including heme-binding cytochromes P450, rather than core biosynthetic genes

Carotenoid trait-associated genes (TAGs) were identified as genes whose expression levels were significantly associated with TC (nonzero log_2_-fold effect size of TC on transcript abundance; adjusted p < 0.05) after accounting for tissue layer and surrogate variables in the negative binomial model. A total of 4,435 TAGs were associated with TC overall, the top 20 of which are shown in Table 3. Of these, 14 belonged to the *a priori* carotenoid gene set, including *DXS, PSY2, phytoene desaturase (PDS), NCED, violaxanthin de-epoxidase (VDE)*, and ζ*-carotene isomerase (ZISO)* (Fig. 7A). However, these carotenoid biosynthetic genes all had relatively low LFC estimates (|LFC| < 0.5 per ug/g TC). Among these TAGs were 54 *a priori* starch-related genes, including *bidirectional sugar transporter SWEET1* (Manes.14G047700; LFC = 0.77, p = 2.0×10^-4^), *pyruvate kinase* (Manes.09G007300; LFC = -0.53, p = 0.03), and *glucose-1-phosphate adenylyltransferase large subunit 1, chloroplastic* (Manes.18G019650; LFC = -0.54, p = 4.2×10^-4^). The set of 491 TAGs with |LFC| > 0.5 (Table S3) were significantly enriched for 10 GO terms (Fig. 8A), which included “heme binding” (p = 2.5×10^-4^), “monooxygenase activity” (p = 2.5×10^-4^), “hexosyltransferase activity” (p = 0.005), and “lipid binding” (p = 0.008). Among the TAGs related to heme binding, there were 5 genes in the *cytochrome P450 CYP2* family, an additional 4 genes in other *cytochrome P450* subfamilies, and 3 *peroxidase*-related genes.

**Figure 7.**
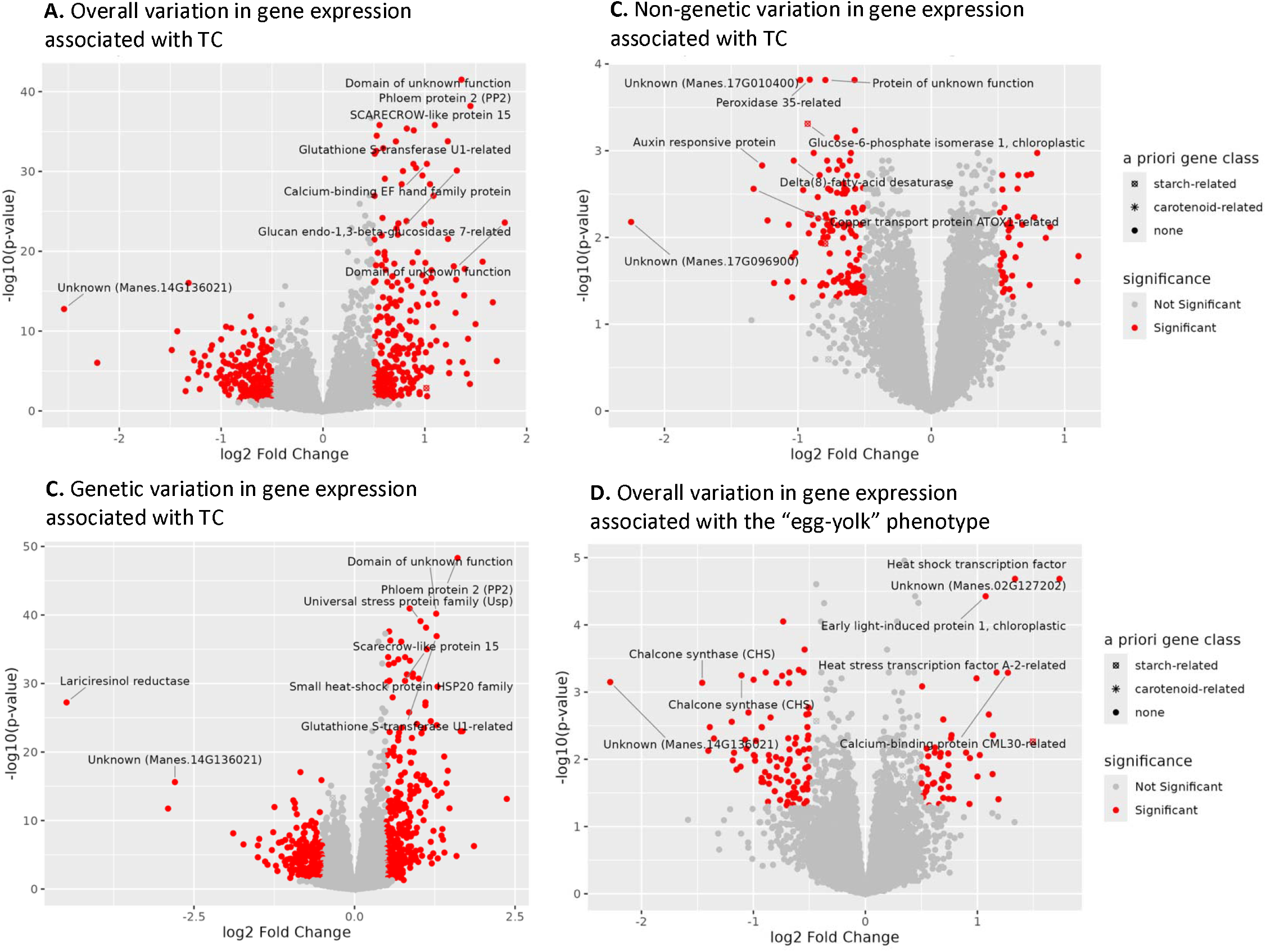
Volcano plots showing the log_2_ fold-change estimates and -log_10_ adjusted p-values for associations between carotenoid traits and components of gene expression: overall variation associated with total carotenoids (TC) (A), genetic variation associated with TC (B), non-genetic variation associated with TC, (C), and overall variation associated with the egg-yolk phenotype (main effect of the difference in TC between inner and outer root layers plus its interaction with tissue layer) (D). Genes in red are considered significant TAGs with adjusted p-value < 0.05 and absolute log_2_ fold-change > 0.5. Genes in the *a priori* carotenoid or starch gene sets are annotated with an asterisk or square, respectively. The eight strongest TAGs in each panel are labeled with their gene/locus name.

**Figure 8.**
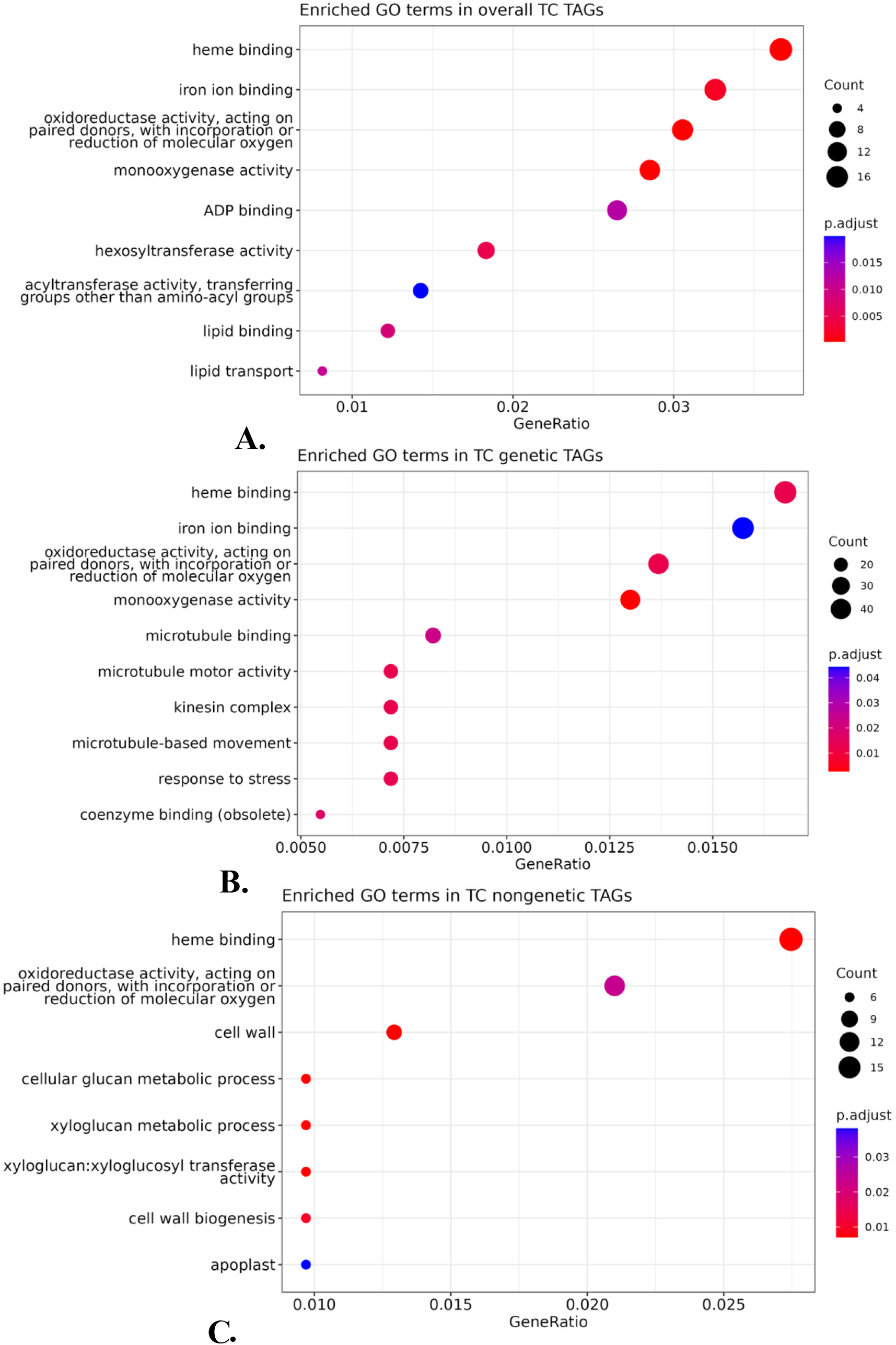
Dotplots of enriched GO terms among genes significantly associated with total carotenoids (TAGs). Panels show TAGs identified using overall variation (A), the genetic component (B), and the non-genetic component (C) of transcript abundance. The x-axis gives the proportion of TAGs annotated with each GO term. Dot size represents the number of TAGs annotated with a given GO term and dot color represents the adjusted p-value from the enrichment test relative to all expressed genes.

Transcriptional variation associated with TC was further partitioned into genetic and non-genetic TAGs. Genetic TAGs were defined as genes associated with TC after accounting for non-genetic experimental covariates, and therefore may reflect transcriptional differences between genotypes with high and low TC. Non-genetic TAGs were defined as genes associated with TC after accounting for genotype-associated differences in transcript abundance, therefore reflecting within-genotype physiological or developmental processes associated with carotenoid accumulation. Analyses of these TAG sets separately can therefore provide insight into whether carotenoid accumulation triggers (or responds to) physiological processes such as differences in carbon partitioning or stress responses, and whether higher expression of carotenoid biosynthetic genes or their regulators drives higher carotenoid accumulation in high TC genotypes.

There were 5,846 genetic TAGs associated with TC, with 523 passing the |LFC| > 0.5 threshold (Fig. 7B; Table S3). Seven GO terms were significantly enriched (Fig. 8B), which mostly overlapped with GO terms enriched in the overall TC TAG set (Fig. 8A). These TAGs included two genes from the *a priori* carotenoid gene set, *NCED* (Manes.03G083500; LFC = 0.53, p = 3.1×10^-4^) and *HMG-CoA reductase* (Manes.01G157500; LFC = -0.81, p = 2.5×10^-3^), and one gene from the *a priori* starch gene set, *glucose-1-phosphate adenylyltransferase large subunit 2, chloroplastic* (Manes.18G019650; LFC = - 0.51, p = 2.2×10^-5^).

There were 1,239 non-genetic TAGs associated with TC, including 161 that passed the |LFC| > 0.5 threshold (Fig. 7C; Table S3). The non-genetic TAGs were significantly enriched for 15 GO terms, including “carbohydrate metabolic process” (p = 4.6×10^-3^), “cellular glucan metabolic process” (p = 1.3×10^-6^), “DNA-binding transcription factor activity” (p = 0.017), “hydrolase activity: (p = 1.8×10^-3^); and “cell wall” (p = 3.0 × 10^-6^) (Fig. 8C). There were 4 *a priori* starch genes among these TAGs: *glucose 6-phosphate isomerase 1, chloroplastic* (Manes.02G146400; LFC = -0.93, p = 4.9×10^-4^), *acid beta-fructofuranosidase 3* (Manes.01G076500; LFC = -0.81, p = 0.01), *pyruvate decarboxylase 2* (Manes.08G094400; LFC = -0.80, p = 0.01), and *hexokinase-3-related* (Manes.06G161400; LFC = -0.58, p = 0.02). There were no *a priori* carotenoid genes among the non-genetic TAGs that passed the LFC threshold.

Spatial variation in carotenoid accumulation was similarly associated mostly with regulatory genes. There were 626 TAGs significantly associated with the egg-yolk phenotype (egg-yolk main effect + egg-yolk/layer interaction), of which 142 passed the LFC threshold (Fig. 7D; Table S3). The top egg-yolk TAGs were all related to regulatory activity (e.g., transcription factors, tRNA charging, protein ubiquitination and binding, and signal transduction) (Table 4), but no GO terms were significantly enriched. There were no statistically significant TAGs for only the egg-yolk/layer interaction effect without the main effect.

### Gene-metabolite networks constructed from estimates of genetic and non-genetic trait variation

Three unsigned, weighted gene/metabolite co-expression networks were constructed using log_2_-fold estimates of the overall phenotypic, genetic, and non-genetic components of gene expression/metabolite abundance. The networks included 20,450 gene/metabolite nodes, including 240 genes and 5 metabolites classified *a priori* as starch-related nodes, and 63 genes and 14 metabolites classified *a priori* as carotenoid-related nodes.

The three networks differed in global topology. The overall network had 24 assigned co-expression modules, with 19% of gene/metabolite nodes remaining in the unassigned/grey module. In comparison, the genetic network had a more granular structure, with 102 assigned modules and only 2% of nodes unassigned. The non-genetic network was less modular, with 16 assigned modules, 36% of nodes unassigned, and a large “turquoise” module containing 6,654 nodes. Since the non-genetic network had a larger unassigned component and fewer, broad modules, module-level interpretation therefore focused on the overall and genetic networks.

### Carotenoid metabolite modules differ between overall and genetic networks

In the overall gene/metabolite network, most of the carotenoid metabolites (phytoene, phytofluene, ζ-carotene, β-carotene, and TC) clustered in the “purple” module, apart from any carotenoid-related genes (Fig. 9; Table S4). The most highly connected hub gene of this module was Manes.09G098760, annotated as a *Universal stress protein (Usp) family* gene, which was strongly positively correlated with TC (r = 0.64, p < 2.2 × 10^-16^). “Purple” module genes with the strongest topological overlap with carotenoids are listed in Table 2. No GO terms were significantly enriched among the 290 genes.

**Figure 9.**
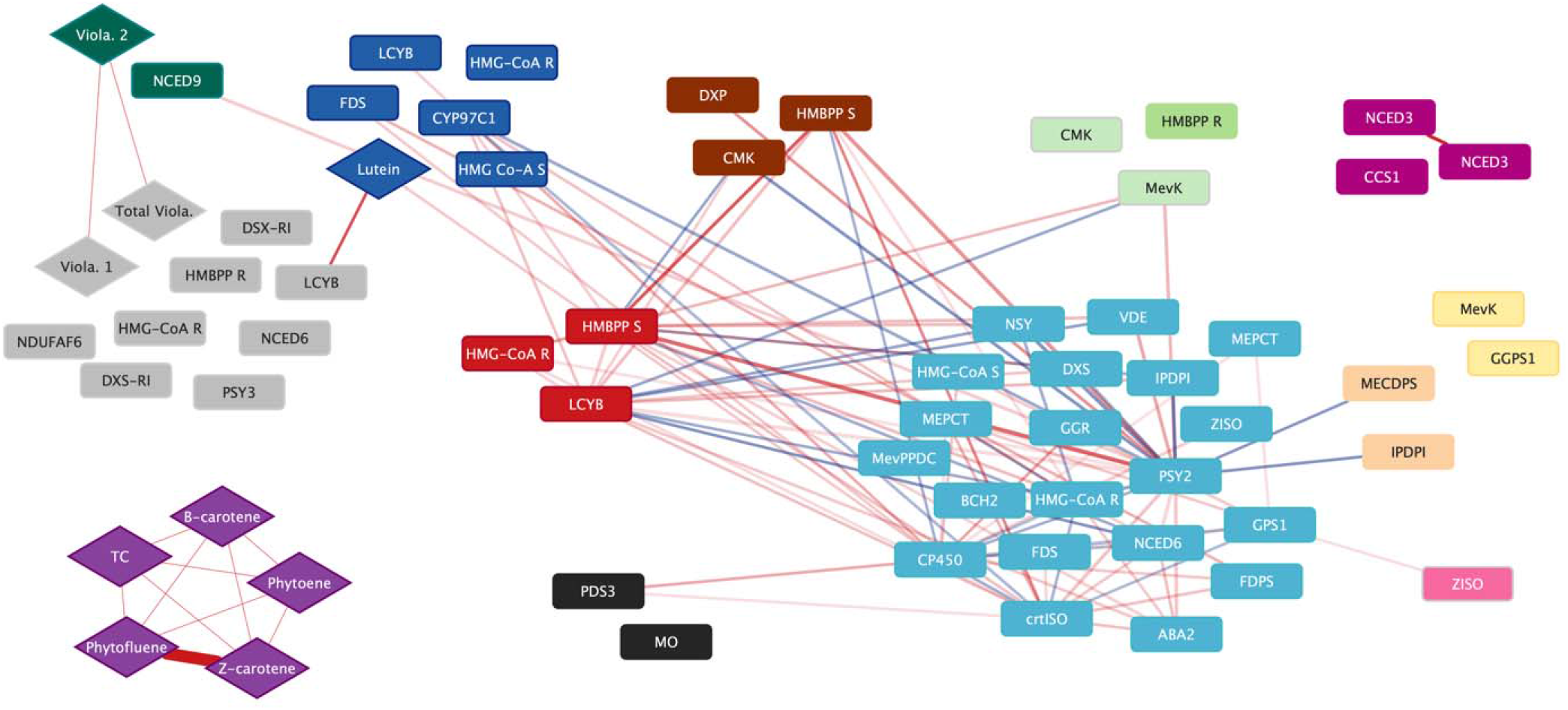
Distribution of carotenoid-related genes/metabolites across co-expression modules in the overall network showing that the major carotenoid compounds in the purple module cluster separately from the majority of the carotenoid biosynthesis genes in the turquoise module. Carotenoid metabolites are represented by diamonds and genes are represented by rectangles. The shape color represents the co-expression module name assigned by WGCNA. Lines represent network edges, with line thickness representing the strength of the topological overlap measure and color representing the direction of the correlation (red = positive, blue = negative). Only edges with weights > 0.008 are shown for clarity. Figure created with Cytoscape. Abbreviations are listed in Table S5.

In the genetic network, carotenoid metabolites showed partial clustering with several carotenoid-related genes. Most carotenoid metabolites clustered in the “black” module along with *PSY2* (Manes.01G124200), *phytoene desaturase* (Manes.05G193700), *4-hydroxy-3-methylbut-2-enyl diphosphate synthase* in the MEP pathway (Manes.11G144700), and *violaxanthin de-epoxidase* (Manes.12G144300). Therefore, genetic differences among genotypes in carotenoid content showed partial co-expression with carotenoid pathway genes, a pattern not observed in the overall network. However, the effect sizes of these genes on TC were small (|LFC| < 0.5), and the “black” module was not defined by isoprenoid synthesis overall. Among the 587 genes in the “black” module, no GO terms were significantly enriched after p-value correction. The hub gene of this module was Manes.03G111200, annotated as a *UDP-glycosyltransferase*, which was strongly positively correlated with TC (r = 0.84, p = 2.27×10^-7^), but negatively correlated with the expression of *PSY2* (r = -0.64, p = 0.0008) and *phytoene desaturase* (r = -0.54, p = 0.007). There were nine carbohydrate-related genes that also clustered with carotenoids in the “black” genetic module, among them *isoamylase (chloroplastic), alpha-amylase*, and *pyruvate kinase* (Table S4).

Among carotenoid metabolites in the overall network, lutein had the highest overall connectivity (k = 73). ζ-carotene and percent *cis-*carotenes also had relatively high connectivities (k=56 and 54 respectively), compared with TC at (k=16) and β-carotene (k=14), indicating that *cis-*carotenes and carotenoid composition overall were more connected to transcript abundance than total carotenoid concentration alone. The distribution of connectivity values for all carotenoids and carbohydrate genes is shown in Fig. S9.

Genes in the ABA biosynthesis pathway, which have been hypothesized to be involved in the TC/DM trade-off, also clustered separately from the carotenoid metabolites in both the overall and genetic networks. For example, in the genetic network, *NCEDs* (Manes.03G083500 and Manes.15G102000) clustered in the “yellow” module along with other phytohormone-related genes, including *jasmonic acid-amino synthetase* (Manes.01G248400) and a *brassinosteroid-responsive RING-H2 zinc finger protein* (Manes.01G254500). No measured carotenoid metabolites were present in the “yellow” module.

### Carotenoid and starch pathway genes do not exhibit significant topological overlap

The permutation test of the overall TOM_G+E_ showed that the carotenoid gene set did not have higher overall total topological overlap with the starch gene set than did sets of random genes with similar connectivity to carotenoids (p=0.56) (Fig. 10). The permutations of TOM_G_ and TOM_E_ also did not show higher genetic or non-genetic topological overlap between carotenoid and starch gene sets (p=0.73 and p=0.94, respectively). The permutation analysis was repeated on an overall network constructed only with the subset of year 2 data, which exhibited a stronger negative phenotypic correlation between TC and DM than in year 1, with similarly statistically insignificant results (p=0.42). In addition, none of the carbohydrate metabolites (glucose, sucrose, starch (mg/g dry weight) nor dry matter (% fresh weight) clustered with carotenoid genes or metabolites in the genetic network.

**Figure 10.**
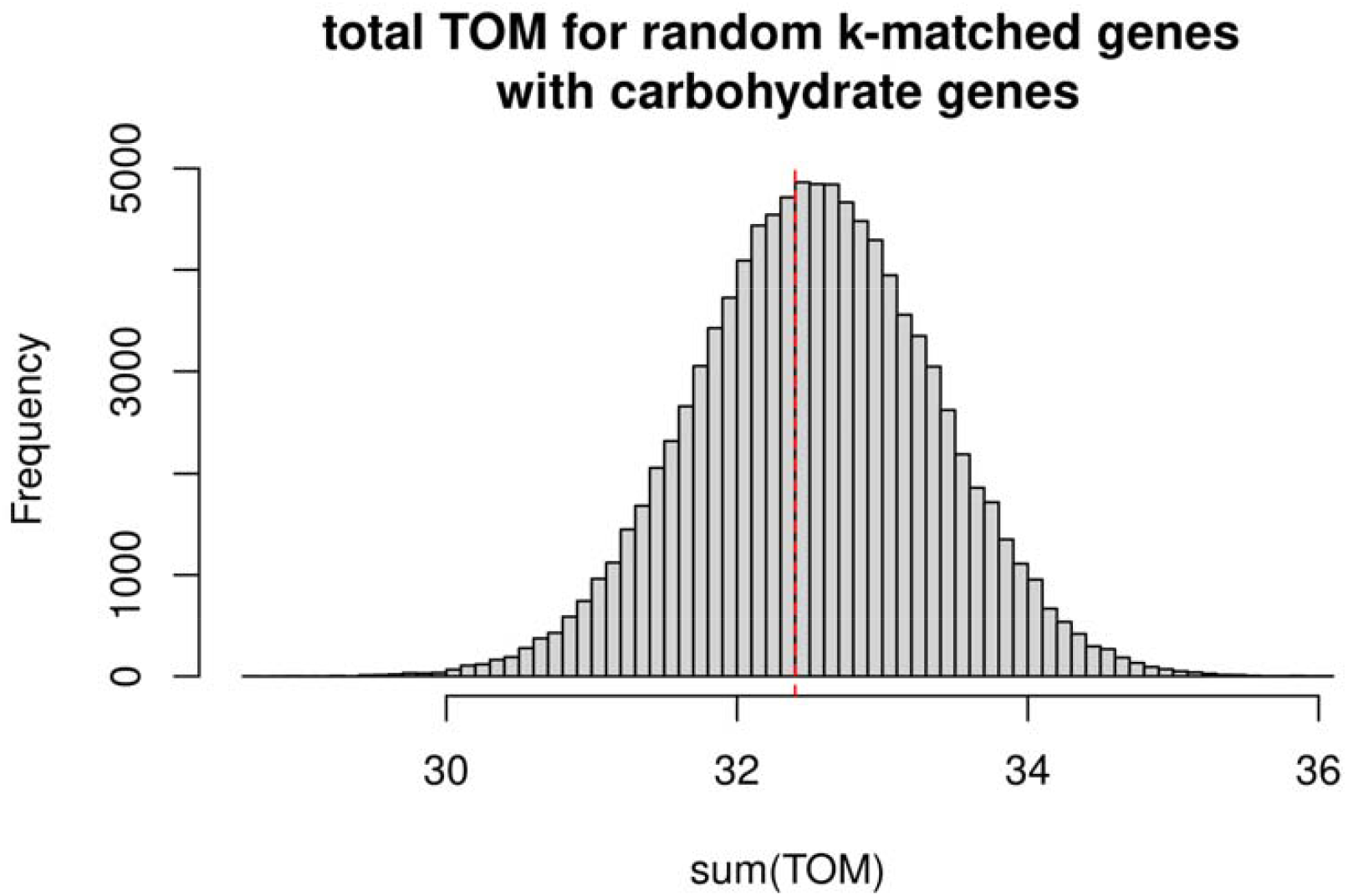
Histogram of the total TOM_G+E_ between a set of random genes (matching the size and connectivity of the carotenoid gene set) and carbohydrate-related genes across 100,000 permutations. The vertical red dashed line indicates the observed total TOM_G+E_ between the carotenoid and carbohydrate gene sets.

While there was no overall topological overlap of the two gene sets, there were some strong associations between individual carotenoid metabolites and starch genes. The genes/metabolites with the strongest topological overlap between carotenoid and carbohydrate classes in the overall network are listed in Table 1 and illustrated in Fig. 11. The starch-related genes most strongly connected to TC in the overall network were *pyruvate kinase* (Manes.07G056100), *pectin lyase* (Manes.13G028600), and *F23N19*.*12*, a predicted pectinesterase inhibitor (Manes.02G034900).

**Table 1.** Summary of the 20 strongest topological overlap measures (TOM_G+E_) between starch-related genes and carotenoid metabolites in the overall network. The Pearson correlations *r* between the overall, genetic, and non-genetic estimates of the carbohydrate genes and carotenoid metabolites are listed.

| Gene Name | Carotenoid | TOM | Gene Description | r_overall | r_genetic | r_nongene |
| --- | --- | --- | --- | --- | --- | --- |
| Manes.07G056100 | Zeta-Carotene | 0.0125 | PYRUVATE KINASE // SUBFAMILY NOT NAMED | -0.48 | -0.63 | -0 |
| Manes.02G034900 | Zeta-Carotene | 0.0095 | F23N19.12 | 0.39 | 0.75 | -0 |
| Manes.02G034900 | Total Carotenoids | 0.0085 | F23N19.12 | 0.47 | 0.73 | 0 |
| Manes.07G056100 | Phytofluene | 0.0082 | PYRUVATE KINASE // SUBFAMILY NOT NAMED | -0.42 | -0.52 | -0 |
| Manes.02G034900 | Beta-Carotene | 0.0080 | F23N19.12 | 0.46 | 0.71 | 0 |
| Manes.13G028600 | Phytofluene | 0.0080 | PECTIN LYASE-LIKE PROTEIN | -0.45 | -0.44 | -0 |
| Manes.13G028600 | Phytoene | 0.0078 | PECTIN LYASE-LIKE PROTEIN | -0.44 | -0.42 | -0 |
| Manes.08G108800 | Phytofluene | 0.0076 | PECTIN LYASE-LIKE PROTEIN | 0.47 | 0.46 | 0 |
| Manes.08G108800 | Phytoene | 0.0075 | PECTIN LYASE-LIKE PROTEIN | 0.48 | 0.45 | 0 |
| Manes.07G056100 | Phytoene | 0.0075 | PYRUVATE KINASE // SUBFAMILY NOT NAMED | -0.39 | -0.52 | -0 |
| Manes.13G028600 | Zeta-Carotene | 0.0075 | PECTIN LYASE-LIKE PROTEIN | -0.44 | -0.40 | -0 |
| Manes.07G056100 | Beta-Carotene | 0.0074 | PYRUVATE KINASE // SUBFAMILY NOT NAMED | -0.40 | -0.52 | -0 |
| Manes.07G056100 | Total Carotenoids | 0.0072 | PYRUVATE KINASE // SUBFAMILY NOT NAMED | -0.42 | -0.54 | -0 |
| Manes.08G108800 | Zeta-Carotene | 0.0066 | PECTIN LYASE-LIKE PROTEIN | 0.45 | 0.46 | 0 |
| Manes.13G028600 | Beta-Carotene | 0.0055 | PECTIN LYASE-LIKE PROTEIN | -0.42 | -0.43 | -0 |
| Manes.13G028600 | Total Carotenoids | 0.0055 | PECTIN LYASE-LIKE PROTEIN | -0.46 | -0.44 | -0 |
| Manes.02G034900 | Phytofluene | 0.0053 | F23N19.12 | 0.34 | 0.64 | -0 |
| Manes.02G034900 | Phytoene | 0.0050 | F23N19.12 | 0.36 | 0.61 | 0 |
| Manes.02G110200 | Zeta-Carotene | 0.0038 | ACONITASE // ACONITATE HYDRATASE 1 | -0.34 | -0.44 | -0 |

**Figure 11.**
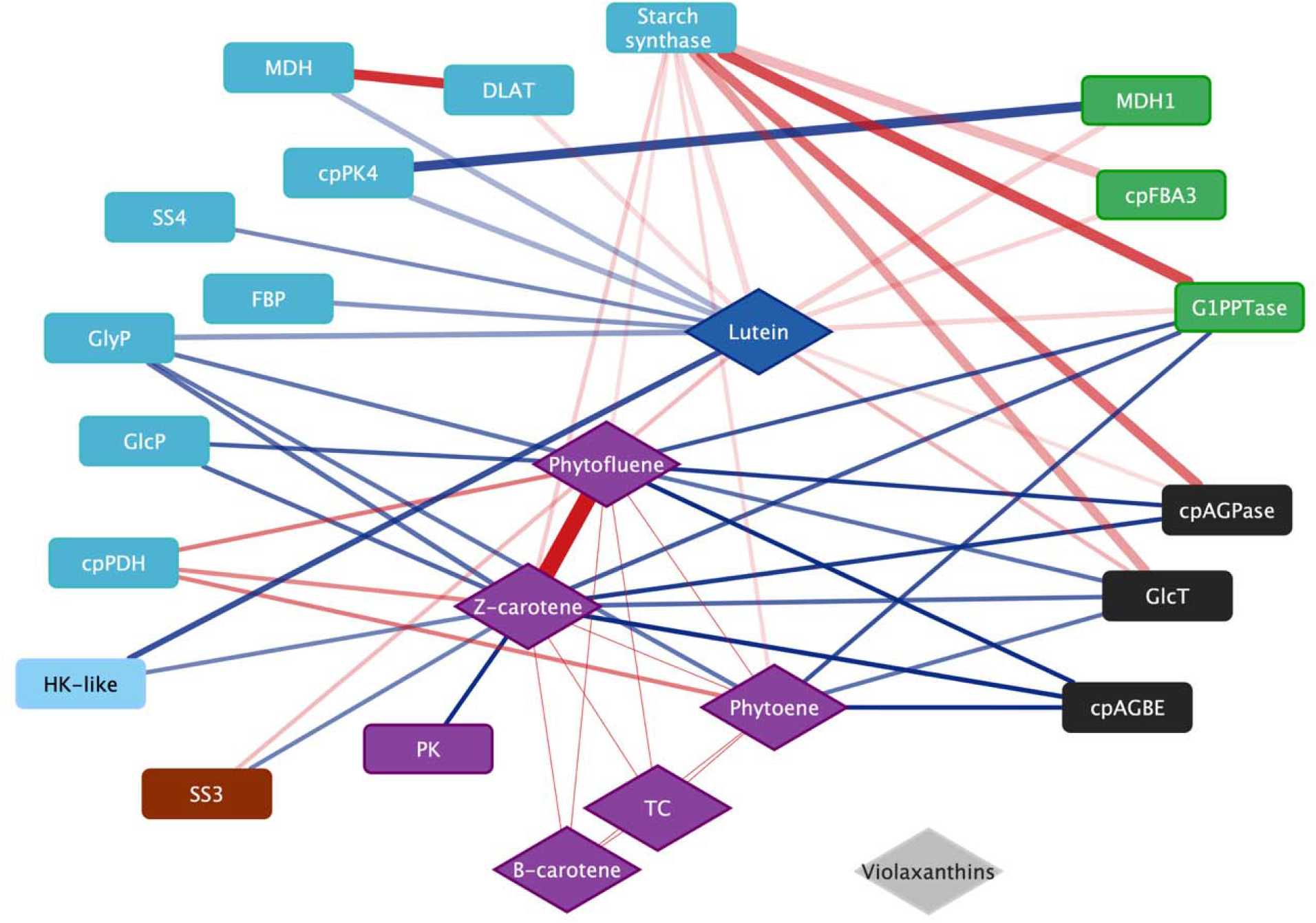
Visualization of the strongest connections between starch-related genes and carotenoid metabolites in the overall co-expression network. Rectangles represent genes and diamonds represent metabolites, and the color of the shape represents the co-expression module they belong to. Lines represent network edges, with line thickness representing the strength of the topological overlap measure and color representing the direction of the correlation (red = positive, blue = negative). Only edges with TOM > 0.011 are shown for clarity. Figure created with Cytoscape. Gene abbreviations are listed in Table S5.

## Discussion

While the carotenoid synthesis pathway is well-characterized, the regulation of carotenoid accumulation in roots and its interaction with carbohydrate metabolism are not well-understood beyond the major effect of the *PSY2* variant (Welsch et al., 2010; Rabbi et al., 2017). Given the persistent negative genetic correlation between TC and DM in cassava roots and other starchy crops, we hypothesized that carotenoid and starch metabolism would be at least partially transcriptionally co-regulated. While the correlation between TC and DM has primarily been observed across genetic diversity, under the hypothesis of an inherent physiological trade-off between carotenoids and starch in amyloplasts, we expected that the traits would also covary not only across genotypes but also among clonal replicates, tissues, or developmental stages of same genotype. Therefore, we aimed to characterize the transcriptional regulation of carotenoids across both genetic diversity and non-genetic phenotypic variation (root tissue layers and developmental stages) and its relationship with starch metabolism.

Genetic and phenotypic correlations across two years of greenhouse trials showed that the negative relationship between TC and DM was weaker and more developmentally influenced than expected based on previous studies. The correlation differed in direction across years and tissue layers (Fig. 3) and became more negative over developmental time during storage root bulking (Fig. 4). Therefore, the strength and direction of the TC/DM association appear to depend on developmental stage and tissue rather than representing a fixed constraint. Across years, a significant effect of TC on DM after accounting for genotype indicated that both genetic and non-genetic factors influencing the traits contributed to their covariation, supporting the hypothesis of a physiological trade-off between the traits. The relationship between carotenoids and DM seemed to be driven primarily by *cis-*carotenes (phytoene, phytofluene, and ζ-carotene) despite their relatively minor contributions to TC, since only *cis-*carotenes were significantly associated with DM when TC was decomposed into upstream *cis-*carotenes and downstream carotenoids in the linear model.

To better understand the physiological basis of the relationship between carotenoids and carbohydrates, we used two complementary approaches: trait-associated gene (TAG) analysis, identifying associations between individual genes and carotenoid metabolites, and gene-metabolite network analysis, which used topological overlap measures (TOMs) to identify shared correlation structure among many genes and metabolites. In both approaches, variation in gene expression and metabolite abundance was parsed into genetic and non-genetic components using generalized linear models, allowing us to distinguish genotype-level associations from physiological relationships occurring within tissues of the same genotype.

To test the hypothesis of transcriptional regulation between the carotenoid and starch pathways, the total topological overlap between all *a priori* carotenoid- and starch-related genes in the gene-metabolite network was compared to a distribution of random pairs of starch-related genes with other genes that had similar overall connectivity values to the carotenoid-related gene set. The permutation test showed that the topological overlap between carotenoid and starch genes was no higher than that of random gene sets, indicating that the pathways are not coordinated directly at the transcriptional level (Fig. 10). In addition, the co-expression modules containing carotenoid metabolites were not strongly defined by carbohydrate metabolic processes.

Although carotenoid and starch pathway gene sets did not show significant topological overlap or evidence of coordinated transcriptional regulation, individual associations between carotenoid metabolites and carbohydrate-related genes suggested possible indirect links. Among the strongest carotenoid/carbohydrate node interactions in the overall network were associations between *cis*-carotenes and lutein with carbohydrate-related genes (Table 1; Fig. 11). The *cis-*carotenes, particularly ζ-carotene, had strong topological overlap with *pyruvate kinase*, which also clustered with carotenoid metabolites in both the overall and genetic networks (Table 1; Table S4). Since *pyruvate kinase* is a key regulatory step in glycolysis, which supplies pyruvate and glyceraldehyde-3-phosphate to the MEP pathway upstream of carotenoid synthesis in the plastid, this association is consistent with a hypothesized link between amyloplast carbon metabolism and carotenoid precursor supply (Villwock et al., 2024). Multiple *pectin lyase-*like genes were also strongly associated with carotenoids, especially *cis-*carotenes but also TC (Table 1). Since pectin is a major component of cell walls that provides rigidity and contributes to cell adhesion, increased expression of *pectin lyase* genes suggest cell wall remodeling through pectin degradation (Anderson and Pelloux, 2026). In addition, the enrichment of “cell wall” and “xyloglucan metabolic process” ontology terms among non-genetic carotenoid TAGs further suggests that within-genotype carotenoid variation was associated with cell wall polysaccharide metabolism, possibly reflecting developmental processes during storage root bulking (Fig. 8C). These results are consistent with the transcriptomic and metabolomic shifts toward cell wall polysaccharide metabolism in high-carotenoid, low-starch cassava genotypes reported by Gutschker et al. (2024). Together, these results support an indirect physiological relationship of carotenoids with broader carbohydrate metabolism and root tissue development, rather than a direct transcriptional relationship with the starch synthesis pathway.

The network analysis also identified potential regulators of carotenoid accumulation independent of carbohydrate metabolism in cassava roots. In the overall network, carotenoid metabolites were more strongly connected to regulatory genes with roles in stress response and signaling (Table 2) than with the core carotenoid biosynthetic genes themselves, suggesting that the carotenoid regulation may occur through post-transcriptional or indirect mechanisms. Transcriptional regulation of carotenoid content by genes outside of the main biosynthetic pathway has also been reported previously in cassava and in carrot (Coe et al., 2023; Olayide et al., 2023), unlike in maize and other crops where most carotenoid variation is driven directly by the allelic variation of the biosynthetic genes or their transcriptional regulation (Baseggio et al., 2020; Diepenbrock et al., 2021; Hershberger et al., 2022). However, in the genetic network, carotenoid metabolites did cluster with several core carotenoid and plastid isoprenoid pathway genes, including *PSY2* and *phytoene desaturase*, suggesting that there is some level of differential transcription of carotenoid pathway genes between genotypes. The hub gene of this module, Manes.03G111200, encodes a UDP-glycosyltransferase, a class of enzymes involved in glycosylation-mediated regulation of secondary metabolism and abiotic stress responses in plants (Gharabli et al., 2023). Its previous association with carotenoid content in African cassava further supports it as a candidate carotenoid regulator for functional investigation (Olayide et al., 2023).

**Table 2.** Summary of the 20 strongest topological overlap measures (TOM_G+E_) between TC and genes clustered with it in the overall network “purple” co-expression module. The Pearson correlations *r* between the overall, genetic, and non-genetic estimates of the genes and carotenoid metabolites are listed.

| Gene Name | Description | TOM | overall_r | genetic_r | nongenetic_r |
| --- | --- | --- | --- | --- | --- |
| Manes.08G105600 | F-box domain (F-box) // Phloem protein 2 (PP2) | 0.069 | 0.69 | 0.84 | 0.22 |
| Manes.16G130900 | Enhancer of rudimentary | 0.062 | 0.57 | 0.75 | 0.10 |
| Manes.09G098760 | Universal stress protein family (Usp) | 0.060 | 0.60 | 0.77 | 0.11 |
| Manes.09G098800 | GLUTATHIONE S-TRANSFERASE, GST, SUPERFAMILY, GST DOMAIN CONTAINING // GLUTATHIONE S-TRANSFERASE U1-RELATED | 0.056 | 0.59 | 0.78 | 0.14 |
| Manes.07G104000 | TETRASPANIN-5-RELATED | 0.055 | 0.59 | 0.73 | 0.09 |
| Manes.04G074700 | Senescence-associated protein (Senescence) | 0.051 | 0.60 | 0.78 | 0.25 |
| Manes.02G033500 | F5O11.5 | 0.045 | 0.60 | 0.79 | 0.17 |
| Manes.18G101100 | PROPROTEIN CONVERTASE SUBTILISIN/KEXIN // SERINE PROTEASE-LIKE PROTEIN-RELATED | 0.045 | 0.52 | 0.74 | 0.08 |
| Manes.05G160600 | GLUCAN ENDO-1,3-BETA-GLUCOSIDASE 7-RELATED | 0.042 | 0.53 | 0.79 | 0.02 |
| Manes.01G150600 | YMGG-like Gly-zipper (Gly-zipper_YMGG) | 0.041 | 0.55 | 0.78 | 0.09 |
| Manes.14G172800 | BCR-ASSOCIATED PROTEIN, BAP // GENOMIC DNA, CHROMOSOME 3, P1 CLONE: MIG5 | 0.041 | 0.54 | 0.71 | 0.07 |
| Manes.01G022400 |  | 0.040 | 0.62 | 0.70 | 0.14 |
| Manes.15G141200 | Deoxyribonuclease V / Escherichia coli endodeoxyribonuclease V | 0.038 | 0.57 | 0.81 | 0.06 |
| Manes.12G027800 | nucleoredoxin [EC:1.8.1.8] (NXN) | 0.038 | 0.56 | 0.73 | 0.14 |
| Manes.06G110100 | next to BRCA1 gene 1 protein (NBR1) | 0.038 | 0.47 | 0.70 | 0.06 |
| Manes.16G121600 | PHOSPHOLIPASE D // PHOSPHOLIPASE D ALPHA 1-RELATED | 0.037 | 0.56 | 0.67 | 0.09 |
| Manes.10G020100 | SMALL HEAT-SHOCK PROTEIN HSP20 FAMILY // SUBFAMILY NOT NAMED | 0.037 | 0.62 | 0.80 | 0.22 |
| Manes.01G136850 | Pantetheine-phosphate adenylyltransferase / PPAT | 0.037 | 0.57 | 0.80 | 0.08 |
| Manes.13G030800 | KINASE-LIKE PROTEIN | 0.036 | 0.47 | 0.72 | 0.09 |
| Manes.12G029366 | Metal-dependent phosphohydrolase | 0.036 | 0.50 | 0.60 | 0.15 |

**Table 3.** The top 20 genes most significantly associated with total carotenoids (TC) using overall transcript abundance variation. LFC indicates the estimated log_2_-fold change in normalized transcript abundance per standard deviation increase in total carotenoids after accounting for tissue layer and surrogate variables. LFC_SE indicates the standard error of the LFC estimate, and padj indicates the Benjamini-Hochberg false discovery rate adjusted p-value.

| locusName | baseMean | LFC | LFC_SE | padj | Description | GO Term description | GO Term ID |
| --- | --- | --- | --- | --- | --- | --- | --- |
| Manes.11G085200 | 44.44 | 1.36 | 0.09 | 3.18E-42 | Domain of unknown function (DUF4408) |  |  |
| Manes.08G105600 | 255.31 | 1.45 | 0.11 | 6.49E-39 | F-box domain (F-box) // Phloem protein 2 (PP2) | protein binding | GO:0005515 |
| Manes.16G130900 | 438.50 | 0.47 | 0.04 | 1.97E-37 | Enhancer of rudimentary | biosynthetic process; cell cycle; positive regulation of Notch | GO:0007049<br>GO:0045747 |
| Manes.01G150600 | 127.70 | 0.55 | 0.04 | 1.56E-36 | YMGG-like Gly-zipper (Gly-zipper_YMGG) | protein binding; zinc ion binding | GO:0008270 |
| Manes.13G137100 | 276.14 | 1.10 | 0.08 | 1.56E-36 | SCARECROW-LIKE PROTEIN 15; GRAS family transcription factor |  |  |
| Manes.04G074700 | 592.80 | 0.82 | 0.06 | 4.29E-36 | Senescence-associated protein (Senescence) |  |  |
| Manes.08G096900 | 91.26 | 0.89 | 0.07 | 7.20E-36 | DEHYDRATION-RESPONSIVE ELEMENT-BINDING PROTEIN 2A-RELATED | activity; regulation of DNA-templated transcription | GO:0003700<br>GO:0006355 |
| Manes.08G117800 | 2786.92 | 0.53 | 0.04 | 3.34E-35 | Rubber elongation factor protein (REF) (REF) |  |  |
| Manes.09G098800 | 242.51 | 1.23 | 0.10 | 1.68E-34 | GLUTATHIONE S-TRANSFERASE, U1-RELATED | protein binding | GO:0005515 |
| Manes.03G111200 | 119.48 | 0.72 | 0.06 | 1.74E-34 | GLUCOSYL/GLUCURONOSYL TRANSFERASES // SUBFAMILY NOT NAMED | metabolic process; hexosyltransferase activity | GO:0008152<br>GO:0016758 |
| Manes.02G011000 | 1001.95 | 0.59 | 0.05 | 1.25E-33 | // DYW family of nucleic acid deaminases (DYW_deaminase) |  |  |
| Manes.18G101100 | 609.86 | 0.52 | 0.04 | 2.73E-33 | PROPROTEIN CONVERTASE SUBTILISIN/KEXIN // SERINE PROTEASE-LIKE PROTEIN-RELATED |  |  |
| Manes.06G110100 | 1145.34 | 0.51 | 0.04 | 6.02E-33 | Next to BRCA1 gene 1 protein (NBR1) // ubiquitin-associated (UBA)/TS-N domain-containing protein / octicosapeptide/Phox/Bemp1 (PB1) domain-containing protein | protein binding; zinc ion binding | GO:0005515<br>GO:0008270 |
| Manes.08G134300 | 48.63 | 1.02 | 0.08 | 1.14E-31 | calcium-binding EF hand family protein | calcium ion binding | GO:0005509 |
| Manes.12G029366 | 97.19 | 0.89 | 0.07 | 1.15E-31 | Metal-dependent phosphohydrolase |  |  |
| Manes.18G077024 | 151.79 | 0.91 | 0.07 | 3.76E-31 | F-BOX PROTEIN PP2-B1-RELATED | protein binding | GO:0005515 |
| Manes.05G160600 | 33.87 | 1.31 | 0.11 | 7.77E-31 | GLUCAN ENDO-1,3-BETA-GLUCOSIDASE 7-RELATED | glycosyl compounds; carbohydrate metabolic process | GO:0004553<br>GO:0005975 |
| Manes.07G019300 | 6727.04 | 0.79 | 0.06 | 9.08E-31 | UBIQUITIN // POLYUBIQUITIN 4 | protein binding | GO:0005515 |
| Manes.09G098760 | 118.77 | 0.98 | 0.08 | 3.21E-30 | Universal stress protein family (Usp) | response to stress | GO:0006950 |
| Manes.12G027800 | 808.11 | 0.61 | 0.05 | 8.22E-30 | disulfide reductase (NAD(P)H) activity |  |  |

**Table 4.** The top 20 genes most significantly associated with the “egg-yolk” phenotype (quantified as the difference in total carotenoid accumulation between the inner and outer root tissue layers) using overall transcript abundance variation. LFC indicates the estimated log_2_-fold change in normalized transcript abundance per standard deviation increase in total carotenoids after accounting for tissue layer and surrogate variables. LFC_SE indicates the standard error of the LFC estimate, and padj indicates the Benjamini-Hochberg false discovery rate adjusted p-value.

| locusName | baseMean | LFC | LFC_SE | padj | Description | GO Term Description | GO Term ID |
| --- | --- | --- | --- | --- | --- | --- | --- |
| Manes.10G026300 | 379.09 | 0.35 | 0.06 | 1.11E-05 | POLY A POLYMERASE CID PAP -RELATED // POLY(A) RNA POLYMERASE GLD2 | nucleotidyltransferase activity | GO:0016779 |
| Manes.02G127202 | 79.08 | 1.73 | 0.29 | 2.07E-05 | PPR repeat family // Pentatricopeptide repeat domain // DYW family of nucleic acid deaminases (DYW_deaminase) |  |  |
| Manes.05G081700 | 12.52 | 1.34 | 0.23 | 2.07E-05 | HEAT SHOCK TRANSCRIPTION FACTOR // SUBFAMILY NOT NAMED | DNA-binding transcription factor activity; nucleus; regulation of DNA-templated transcription; sequence-specific DNA binding | GO:0003700<br>GO:0005634<br>GO:0006355<br>GO:0043565 |
| Manes.09G003900 | 31.63 | -0.44 | 0.08 | 2.48E-05 | F-box domain (F-box) | protein binding | GO:0005515 |
| Manes.14G023900 | 413.98 | 0.44 | 0.08 | 3.75E-05 | C2 domain (C2) | protein binding | GO:0005515 |
| Manes.15G033100 | 507.00 | 1.07 | 0.19 | 3.75E-05 | UPF0041 BRAIN PROTEIN 44-RELATED // EARLY LIGHT-INDUCED PROTEIN 1, CHLOROPLASTIC-RELATED |  |  |
| Manes.12G062800 | 93.83 | 0.47 | 0.08 | 4.71E-05 | THREONYL-TRNA SYNTHETASE / THREONINE--TRNA LIGASE-RELATED | nucleotide binding; aminoacyl-tRNA ligase activity; threonine-tRNA ligase activity; ATP binding; cytoplasm; tRNA aminoacylation for protein translation; threonyl-tRNA aminoacylation; tRNA aminoacylation | GO:0000166<br>GO:0004812<br>GO:0004829<br>GO:0005524<br>GO:0005737<br>GO:0006418<br>GO:0006435<br>GO:0016876<br>GO:0043039 |
| Manes.04G146600 | 58.58 | -0.37 | 0.07 | 4.76E-05 | VON WILLEBRAND FACTOR, TYPE A DOMAIN CONTAINING // SUBFAMILY NOT NAMED |  |  |
| Manes.01G214200 | 19.11 | -0.73 | 0.13 | 8.91E-05 | Predicted ncRNA |  |  |
| Manes.06G063900 | 65.00 | -0.40 | 0.07 | 8.91E-05 | Predicted ncRNA |  |  |
| Manes.10G087600 | 48.06 | 0.28 | 0.05 | 8.91E-05 | LEUCINE-RICH REPEAT-CONTAINING PROTEIN // SUBFAMILY NOT NAMED | protein binding; signal transduction; ADP binding | GO:0005515<br>GO:0007165<br>GO:0043531 |
| Manes.01G016300 | 398.77 | -0.37 | 0.07 | 1.09E-04 | tyrosyl-tRNA synthetase (YARS, tyrS) | nucleotide binding; aminoacyl-tRNA ligase activity; tyrosine-tRNA ligase activity; ATP binding; tRNA aminoacylation for protein translation | GO:0000166<br>GO:0004812<br>GO:0004831<br>GO:0005524<br>GO:0006418 |
| Manes.02G199300 | 19.23 | -0.54 | 0.10 | 2.33E-04 | FAMILY NOT NAMED // LEUCINE-RICH RECEPTOR-LIKE PROTEIN KINASE | protein kinase activity; protein tyrosine kinase activity; protein binding; ATP binding; protein phosphorylation | GO:0004672<br>GO:0004713<br>GO:0005515<br>GO:0005524<br>GO:0006468 |
| Manes.06G079900 | 191.08 | 0.19 | 0.04 | 2.33E-04 | RING ZINC FINGER PROTEIN // SUBFAMILY NOT NAMED | protein binding; zinc ion binding | GO:0005515<br>GO:0008270 |
| Manes.06G066900 | 23.18 | -0.59 | 0.12 | 4.68E-04 | WD40-repeat-containing domain superfamily |  |  |
| Manes.02G174100 | 633.33 | -0.39 | 0.08 | 5.10E-04 | BIDIRECTIONAL SUGAR TRANSPORTER SWEET1 |  |  |
| Manes.10G113400 | 9.46 | -0.89 | 0.18 | 5.10E-04 | GLUTATHIONE S-TRANSFERASE, GST, SUPERFAMILY, GST DOMAIN CONTAINING // SUBFAMILY NOT NAMED | protein binding | GO:0005515 |
| Manes.11G082579 | 516.50 | -0.55 | 0.11 | 5.10E-04 | Phosphatidylinositol diacylglycerol-lyase / Phosphatidylinositol phospholipase C |  |  |
| Manes.13G068000 | 1626.67 | 0.21 | 0.04 | 5.10E-04 | PEPTIDASE A1 DOMAIN-CONTAINING PROTEIN // Phytpepsin | aspartic-type endopeptidase activity; proteolysis; lipid metabolic process; lysosome; sphingolipid metabolic process | GO:0004190<br>GO:0006508<br>GO:0006629 |
| Manes.14G024400 | 112.48 | 0.44 | 0.09 | 5.10E-04 | WW domain-containing oxidoreductase (WWOX) | metabolic process; oxidoreductase activity | GO:0008152<br>GO:0016491 |

In addition, among overall and genetic TC TAGs (Table S3), the enrichment of GO terms related to heme/iron binding, monooxygenase, and oxidoreductase activity, predominately driven by *peroxidases* and *cytochrome P450* (*CYP*) genes, suggests an important role of heme-dependent redox enzymes in either regulating carotenoid accumulation or redox processes coordinated with carotenoids. CYPs have diverse roles in the synthesis of secondary metabolites, including those from the CYP97 family known to act as carotenoid hydroxylases (Arango et al., 2014; Lautier et al., 2023) and the CYP707 family involved in abscisic acid regulation (Chakraborty et al., 2023). To our knowledge, the *CYP2* family genes implicated here have not been functionally characterized in cassava and would be interesting targets for follow-up study.

Co-expressed gene modules and TAGs associated with the egg-yolk phenotype (enrichment of TC in the inner root tissue layer) suggest that the spatial regulation of carotenoid accumulation is also driven without direct transcriptional regulation of the carotenoid biosynthetic genes. There were no genes from the *a priori* carotenoid gene set among the 625 TAGs associated with the egg-yolk phenotype. Rather, several TAGs were identified as possible regulators of carotenoid accumulation in cassava. *EARLY LIGHT-INDUCED PROTEIN 1, CHLOROPLASTIC-RELATED* (Manes.15G033100) was a plastid-related TAG (LFC = 1.07, p = 3.8×10^-5^) with relevance to carotenoid accumulation. Early light-induced proteins (ELIPs) are involved in photooxidative protection and bind both chlorophyll and carotenoids (Hutin et al., 2003; Guseinova et al., 2004; Liu et al., 2020). *ELIP* expression has been associated with the chloroplast to chromoplast transition in ripening tomato (Bruno and Wetzel, 2004). Since ELIPs have transmembrane domains and bind carotenoids, Giuliano et al. (1993) suggested that ELIP may facilitate carotenoid accumulation in plastid membranes. In addition, *Photosystem II Core Complex Proteins PsbY, chloroplastic* (Manes.13G019200) was significantly negatively associated with the egg-yolk phenotype (LFC = -0.64, p = 0.023). While the function of PsbY as an auxiliary protein in photosystem II is not clear, it may modulate redox states of photosystem II’s cytochrome b_599_ (Plöscher et al., 2009; Plöchinger et al., 2016). While the association of carotenoids and photosynthesis- and light-related genes in below-ground root tissues may seem counterintuitive, the suppression of photosystem-related components may be involved in chromoplast or amylo-chromoplast differentiation. Finally, several stress-signaling transcription factors (e.g., *heat shock transcription factor/*Manes.05G081700, *heat stress transcription factor A-2 related/*Manes.16G081500, and *heat stress transcription factor C-1/*Manes.03G020500) were positively associated with the egg-yolk phenotype. Together, these TAGs suggest that plastid dynamics and stress-responsive signaling may underlie the spatial pattern of enriched carotenoid accumulation in the inner tissue layer of cassava roots.

One limitation of this study is that the cassava plants were grown in a greenhouse setting, with growing conditions and management practices that are atypical for what cassava plants experience in the field in west Africa. Since greenhouse performance cannot be extrapolated to field performance, yield and other agronomic traits of importance for varietal candidates were not measured. Similarly, dry matter content and carotenoid content measured on these plants are not accurate evaluations of these lines as varietal candidates. Even carotenoid content, a trait with relatively high heritability, has been reported to differ significantly between greenhouse-grown plants and clones of the same accessions in field trials. Therefore, only the relative variation and covariation among traits were interpreted. Since we found the TC/DM relationship to be developmentally and environmentally influenced, different field conditions like drought stress would likely perturb these metabolic relationships. Future studies are needed to evaluate the stability of gene-metabolite associations under field conditions.

In summary, this study provides new insight into the regulation of carotenoid accumulation in cassava and its interactions with carbohydrate metabolism by integrating phenotypic, transcriptomic, and targeted metabolomic data across genetic, spatial, and temporal variation. We found that the TC/DM trade-off in cassava roots was not explained by a single genetic constraint. Instead, associations between carotenoids and DM across genetic diversity, developmental stage, and root tissue layer indicated that both genetic and non-genetic physiological variation contribute to the trade-off. Among carotenoid traits, upstream *cis-*carotenes were most strongly associated with DM despite being a minor component of total carotenoids, suggesting that carotenoid composition is more informative than TC alone. However, gene/metabolite network analysis did not support direct transcriptional co-regulation of the core carotenoid and starch pathways, with carotenoid- and starch-related gene sets showing minimal topological overlap. Therefore, the TC/DM trade-off is likely driven post-transcriptionally or mediated by indirect regulatory mechanisms. The strongest associations between carbohydrate-related genes and carotenoids pointed to plastidial carbon metabolism and cell wall carbohydrate remodeling as possible indirect connections between TC and DM. In addition, the gene/metabolite networks showed that regulatory genes rather than core biosynthetic genes were most strongly associated with carotenoid variation overall. Carotenoid-associated genes related to redox metabolism and stress signaling suggest that carotenoid accumulation is either regulated by or coordinated with oxidative stress responses in cassava roots. Candidate carotenoid regulators such as *CYP2* family genes, *ELIP1*, and *UDP-glycosyltransferase* would be interesting targets for future functional validation. Overall, these results cassava breeders may still be able to make progress on both provitamin A content and dry matter, though future research on the biological mechanisms connecting carotenoid and carbohydrate metabolism would further inform selection strategies and the fundamental understanding of carotenoid variation in starchy tissues.

## Abbreviations

ABA: abscisic acid
BLUE: best linear unbiased estimate
CYP: cytochrome P450
DM: dry matter
ELIP: early light-induced protein
GO: gene ontology
LFC: log2-fold change
MAP: months after planting
MEP: methylerythritol phosphate
NCED: *9-cis-epoxycarotenoid dioxygenase*
PC: principal component
PGO: peroxidase glucose oxidase
PSY2: *phytoene synthase 2*
SVA: surrogate variable analysis
TAG: trait-associated gene
TC: total carotenoids
TOM: topological overlap measure
WGCNA: Weighted Gene Co-expression Network Analysis

## Author Contributions

S.S.V: Conceptualization, Data curation, Formal analysis, Investigation, Writing – original draft, Funding Acquisition. K.G.: Investigation, Formal analysis. T.F.: Investigation, Methodology, Data curation, Formal analysis, Writing – review & editing. J.L.: Investigation. A.D.: Investigation. A.W.: Investigation. T.T.: Methodology, Resources, Supervision, Writing – review & editing. M.G.: Methodology, Supervision, Writing – review & editing. J-L. J.: Conceptualization, Methodology, Resources, Supervision, Writing – review & editing, Funding Acquisition.

## Acknowledgements

We thank the Cornell University BRC Bioinformatics Core Facility (RRID:SCR_021757) for providing access to computational resources and Jeff Glaubitz for technical bioinformatics consultation. We thank Jing Wu and the BRC Genomics Core Facility (RRID:SCR_021727) for conducting RNA library prep and sequencing. Lauren Brzozowski, Michelle Okoma, Luciano Rogério Braatz de Andrade, and Leah Treffer assisted with cassava planting and/or harvesting. Ismail Y. Rabbi and Li Li provided constructive feedback throughout. We are grateful to the staff of the Guterman Bioclimatic Laboratory for plant care, especially growers Janet Myrick, Jerrie Hanes, Shannan Sweet, Amy Soriano, Jeffrey Persky, and Scott Anthony. We thank Tim Setter and Peter Hyde for sharing supplies, equipment, and advice on plant care and carbohydrate assay protocols. We thank Mark Sorrells, Gary Bergstrom, Michael Gore, and Ted Thannhauser and their lab staff for providing bench space and lab supplies, freezer space, and equipment use. We are grateful to Xiaowei Li for providing RNA isolation training and Liam Wickes-Do for lab support. We thank Claire Casteel and lab for maintaining and sharing use of the lyophilizer. We thank Sharon Wages and Joanna Norton at the University of Hawaii at Mānoa for providing cassava stem cuttings and seeds, and Peter Kulakow for assisting with the germplasm transfer.

## Data Availability Statement

All R code and datasets are available on GitHub (https://github.com/serenvillwock/TCDM_network_analysis), except the raw RNA sequencing data, which is publicly available at NCBI Sequencing Read Archive under BioProject ID PRJNA1258369 (http://www.ncbi.nlm.nih.gov/bioproject/1258369). Cassava accessions used in this study were contributed to the National Clonal Germplasm Repository at the USDA Subtropical Horticulture Research Station, Miami, Florida.

