## Supplemental Figures S1-S9 for "Gene-metabolite networks reveal physiological trade-offs but not transcriptional co-regulation between carotenoid and dry matter accumulation in cassava *(Manihot esculenta)* roots"

**Supplementary figures**

**Table (a): Waters UPC2 gradient conditions**

| | Time (min) | Flow (ml/min) | % A (Co2) | % B (Methanol) | curve |
|---|---|---|---|---|---|
| 1 | Initial | 1 | 99 | 1 | Initial |
| 2 | 7.5 | 1 | 80 | 20 | 10 |
| 3 | 10 | 1 | 80 | 20 | 6 |
| 4 | 12 | 1 | 80 | 20 | 6 |
| 5 | 15 | 1 | 99 | 0 | 1 |

**Table (b): ABPR pressure gradient settings**

| | Time (min) | Pressure (psi) |
|---|---|---|
| 1 | Initial | 3750 |
| 2 | 7 | 3750 |
| 3 | 10 | 3000 |
| 4 | 12 | 3000 |
| 5 | 14 | 3750 |

**Table (c): Carotenoids optimal wavelengths and elution times by UPC2**

| Compound | wavelength (nm) | Retention Time (min) |
|---|---|---|
| phytoene | 282 | 3.31 |
| phytofluene | 340 | 3.73 |
| zeta carotene | 411 | 4.24 |
| 9 cis beta carotene | 430 | 6.49 |
| trans beta carotene | 435 | 6.62 |
| Violaxanthin 1 | 415 | 10.66 |
| Violaxanthin 2 | 415 | 11.1 |
| Lutein | 430 | 11.3 |
| DIM | 282 | 8.79 |

**Figure (d): Waters Gradient curves**

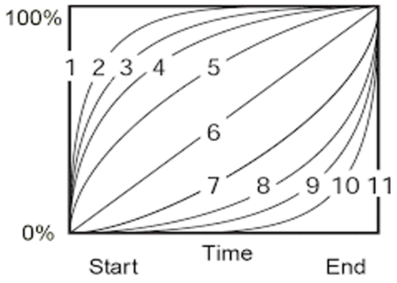


**Figure S1**. Details of the chromatography conditions for carotenoid separation and quantification. A) Waters UPC2 gradient conditions; B) Automated back pressure regulator (ABPR) pressure gradient settings; C) Carotenoids optimal wavelengths and elution times by UPC2; D) Waters Gradient curves.

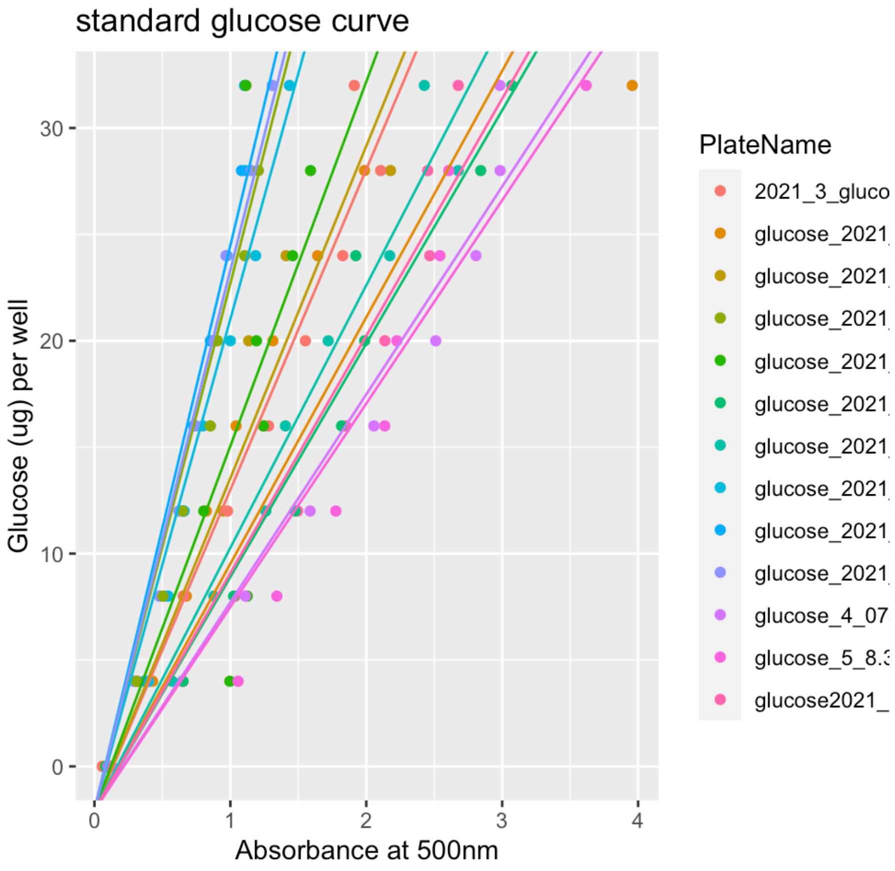


**Figure S2.** Standard glucose curves and the regression line used to predict glucose concentration from absorbance at 500nm after PGO reaction.

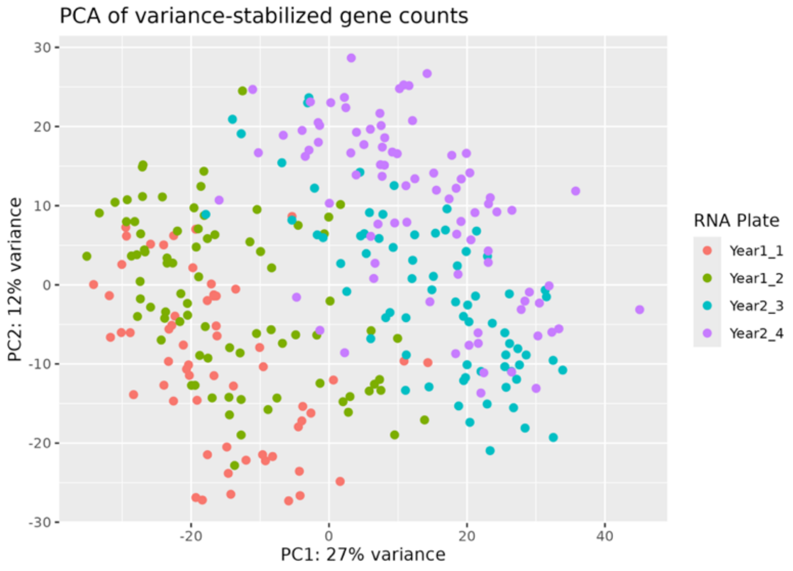


**Figure S3.** Principal component analysis of gene counts after variance-stabilizing transformation with DESeq2 showing stratification of batches by year (different sequencing types) and 96-well plate used for RNA library preparation.

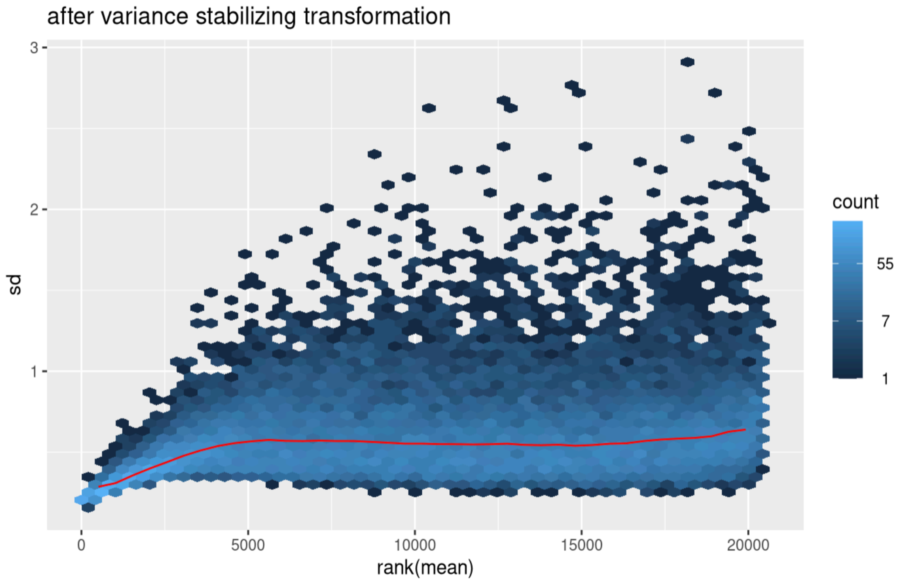


**Figure S4.** Plot of the relationship between gene expression counts mean and variance after the variance-stabilizing transformation accounting for overdispersion.

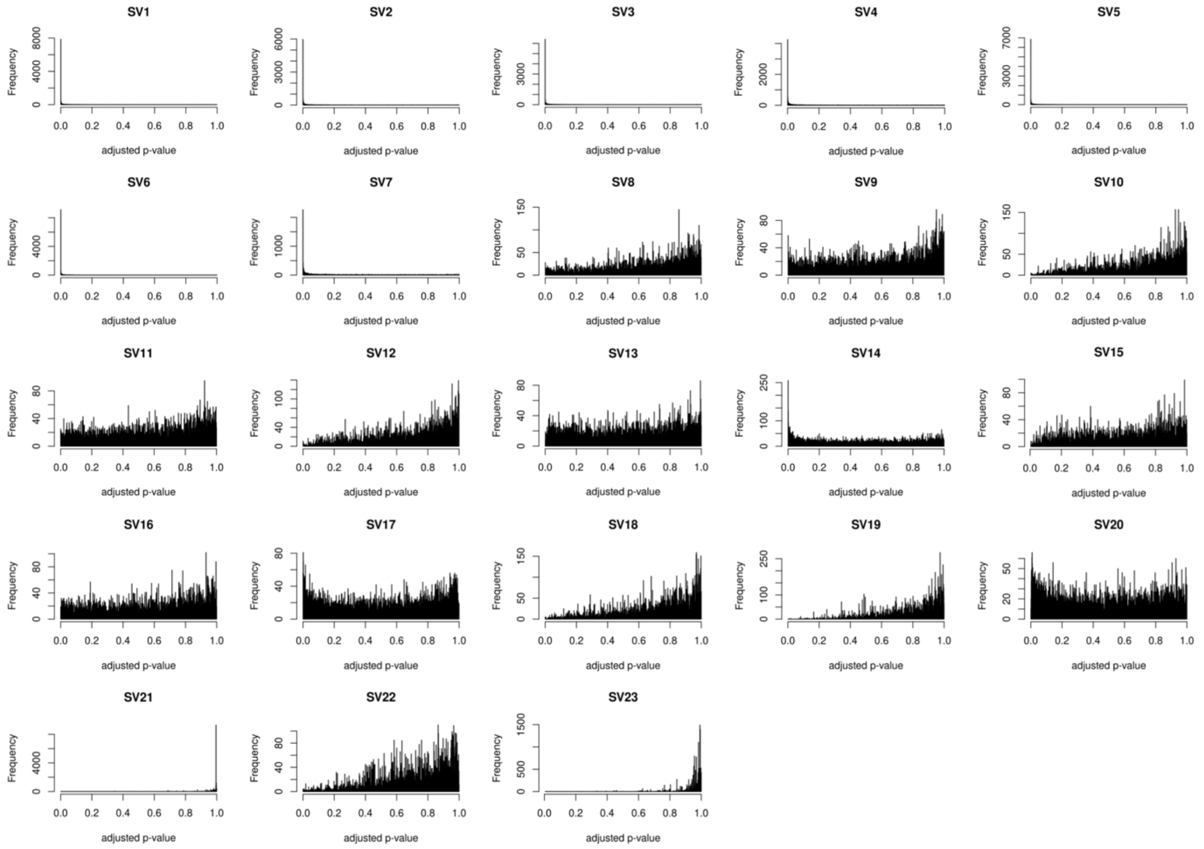


**Figure S5.** Distributions of the p-values from t-tests on estimates of the effect of surrogate variables on gene expression. Surrogate variables were calculated with SVA in R, with the first 23 variables determined to be statistically significant. The first 7 surrogate variables were selected as covariates based on most p-values being near 0, indicating significant associations with most genes, likely reflective of technical factors.

A. Genetic network

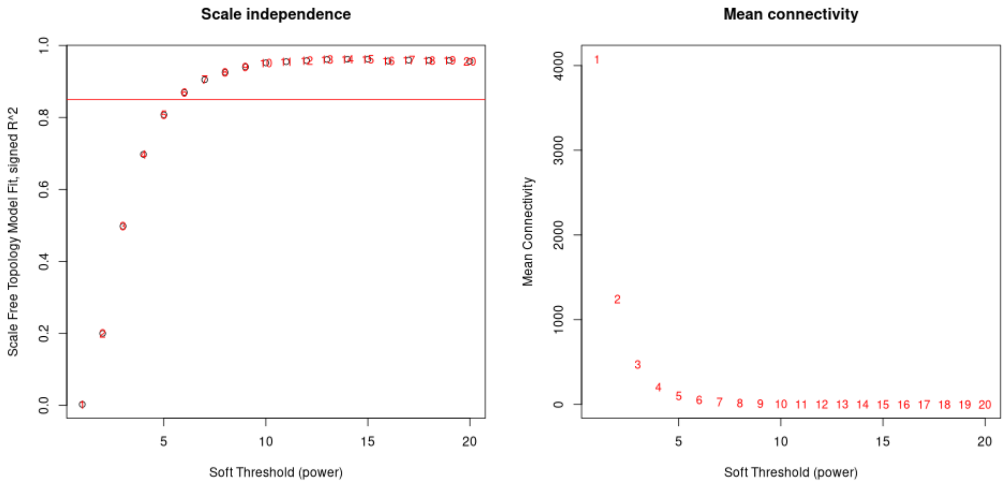


B. Non-genetic network

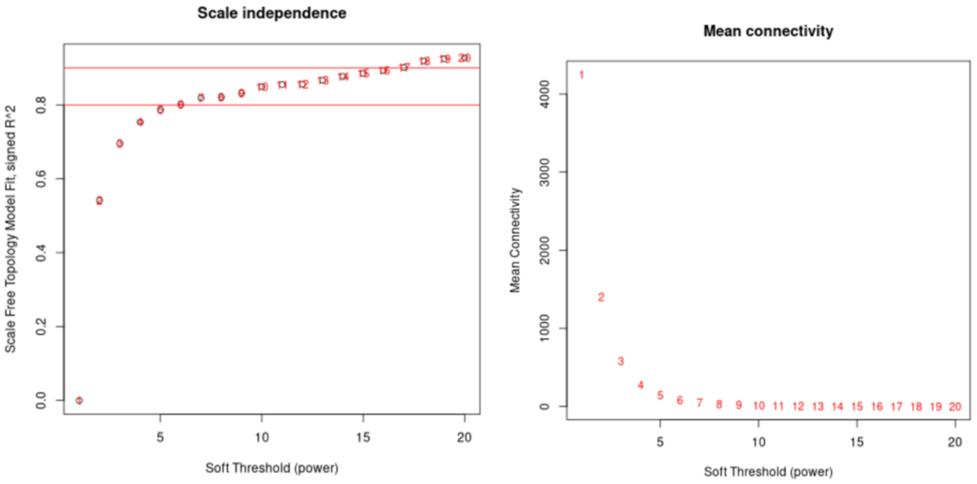


C. Overall network

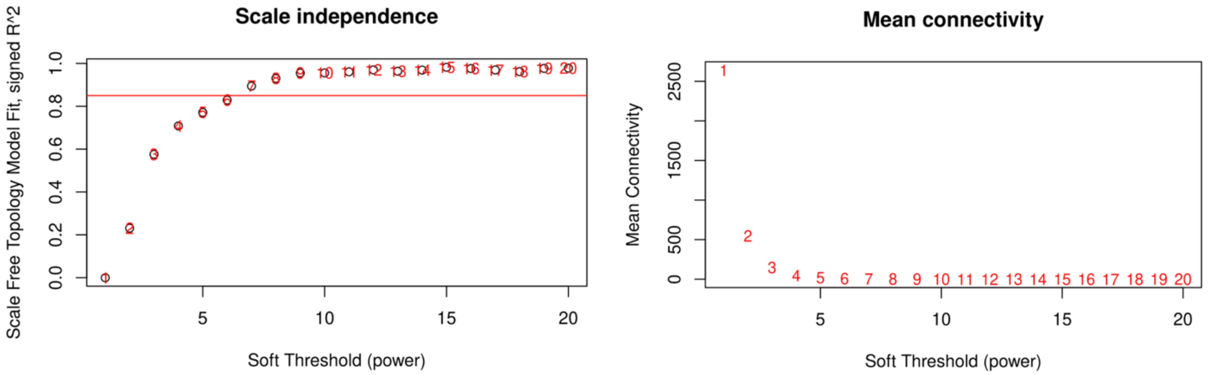


**Figure S6.** Plots of soft thresholding powers tested to determine the lowest power at which the network approximates a scale-free topology ($R^2 > 0.85$) for the A) genetic, B) non-genetic, and C) overall gene/metabolite networks.

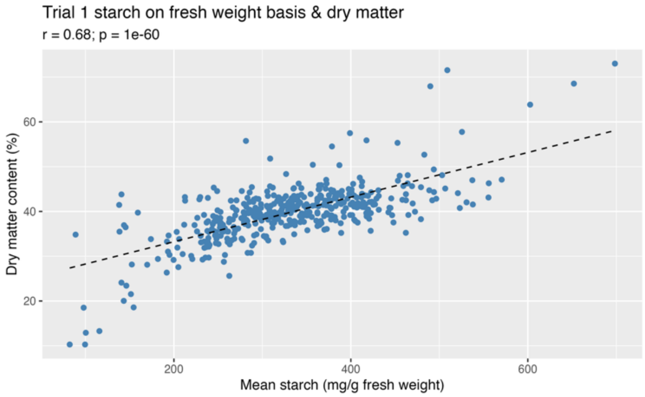


**Figure S7.** Correlation between starch and DM percentage. Each point represents a mean value per sample across two technical replicates. Dashed lines represent the linear regressions of DM on starch.

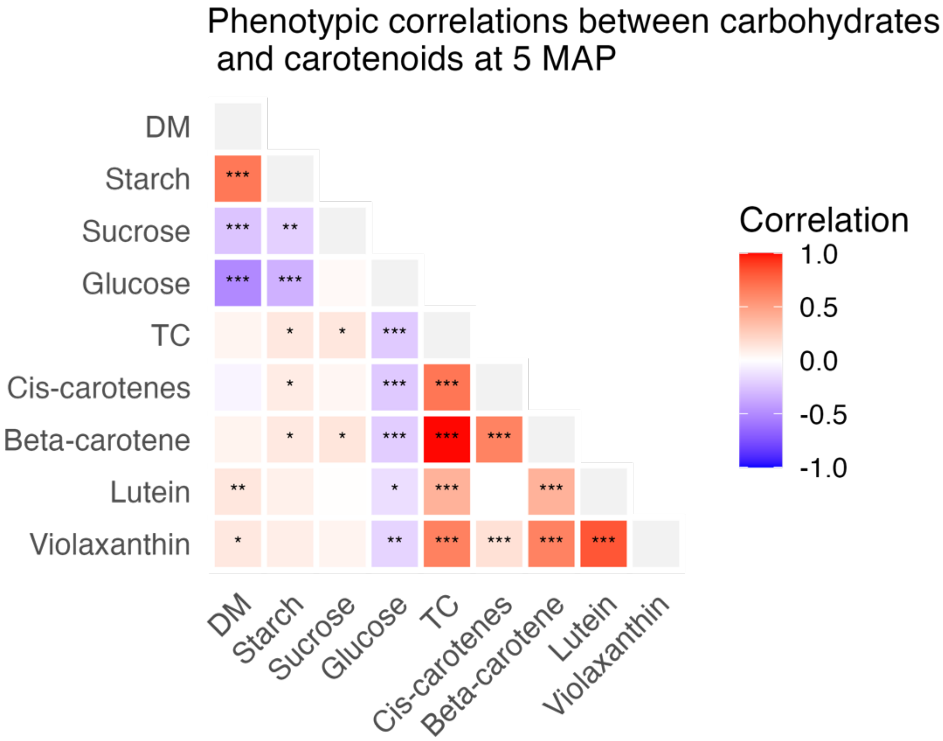


**Figure S8**. Pairwise correlation heatmap of carbohydrates and carotenoids among trial 1 samples harvested at 5 MAP. Starch is expressed on a fresh weight basis for comparison with DM percentage; all other variables are expressed on a dry weight basis. *Cis*-carotenes represents total phytoene, phytofluene, and ζ-carotene; $\beta$-carotene and violaxanthin refer to the total contents of their respective isomers. The color of each cell represents the pairwise Pearson correlation coefficient from -1 (blue) to +1 (red). Asterisks indicate statistically significant correlations: $p < 0.05$ *, $p < 0.01$ **, and $p < 0.001$ ***. Only the lower triangle of the correlation matrix is shown for clarity. TC = total carotenoids; DM = dry matter.

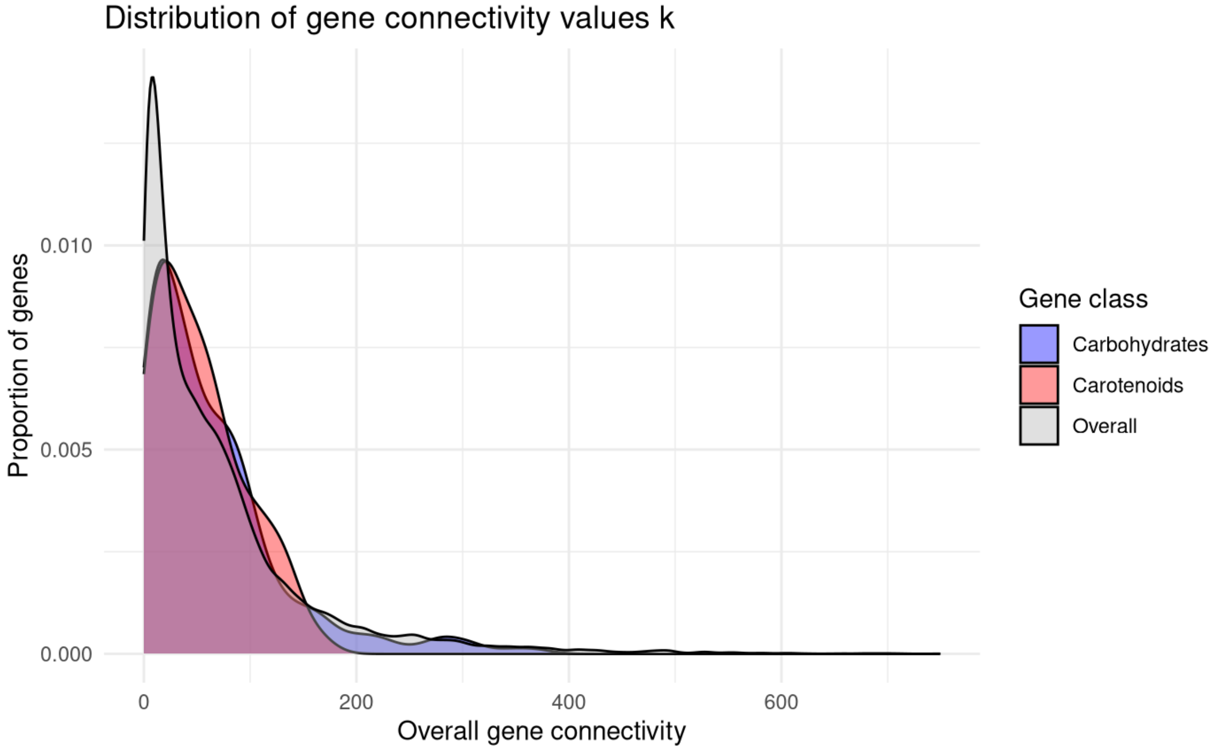


**Figure S9.** Distribution of gene connectivity values *k* across all genes in the overall gene/metabolite network. The y-axis indicates the proportion of genes with a given connectivity value on the x-axis. Genes in the carotenoid gene class are shown in red and genes in the carbohydrates gene class are shown in blue.

**Table S1.** Summary of root tissue samples selected for transcriptomic analysis.

**Table S2.** Genes with GO or Pathway annotations related to carotenoid or carbohydrate metabolism.

**Table S3.** List of trait-associated genes (TAGs) associated with total carotenoids via genetic, non-genetic, and overall components of transcript variation with $\log_2$-fold effect estimates passing the $|LFC| > 0.5$ threshold.

**Table S4.** Module placement of key carotenoid- and carbohydrate-related gene/metabolite nodes in the overall, genetic, and non-genetic co-expression networks.

**Table S5.** Gene/metabolite name abbreviations corresponding to network figures.
